# A canonical input to the head direction compass carries predominantly eye movement signals

**DOI:** 10.64898/2026.10.26.761874

**Authors:** Jeffrey J. Stott, Hui Ho Vanessa Chang, Chenhao Bao, Kechen Zhang, Kathleen E. Cullen, Matthijs A. van der Meer

## Abstract

The mammalian head direction (HD) system is thought to update its internal compass using angular head velocity (AHV) signals relayed from the brainstem. We tested this canonical interpretation by recording from the nucleus prepositus hypoglossi (NPH), a proposed AHV relay known to be important for accurate HD tracking, while simultaneously measuring eye and head movements in mice. Neurons classified as AHV-tuned instead predominantly encoded eye movements: most showed saccade-related bursts or pauses, and apparent AHV tuning was largely explained by eye rather than head motion. Optotagging showed that even NPH neurons projecting directly to the HD circuit predominantly encoded eye movements. Empirically constrained population models reconstructed eye position and gaze direction, but head-direction readouts failed when eye and head motion were dissociated. These findings overturn the canonical view of NPH as a dedicated AHV relay, revealing eye movements as a major, previously unmeasured component of the signals entering the mammalian HD circuit and thereby changing the computational problem that the HD system must solve to maintain a stable representation of heading.

## Introduction

Across species, directional heading is represented by dedicated neural circuits that function as an internal compass. In mammals, head direction (HD) neurons are thought to form a continuous attractor network in which a stable packet of population activity represents current heading and shifts as the animal turns (Sharp et al., 2001a; Taube, 2007; Peyrache et al., 2017; Hulse and Jayaraman, 2020). Related circuit architectures in fruit flies and zebrafish suggest that maintaining a directional representation while continuously updating it from self-motion is a conserved computational problem (Seelig and Jayaraman, 2015; Green et al., 2017; Petrucco et al., 2023). Canonical models of the mammalian HD network propose that its dynamics are governed by two major inputs. Visual signals anchor the compass to environmental landmarks, whereas ascending movement-related signals shift the activity packet as the animal turns (Chen et al., 1994; Goodridge and Taube, 1997; Stackman and Taube, 1997; Bassett et al., 2007; Yoder and Taube, 2009; Clark et al., 2010; Page and Jeffery, 2018). In continuous-attractor models, this movement-related input is explicitly specified as angular head velocity (AHV) (Skaggs et al., 1995; Redish et al., 1996; Zhang, 1996; Song and Wang, 2005; Kim et al., 2017; Laurens and Angelaki, 2018).

Anatomically, the nucleus prepositus hypoglossi (NPH) provides a major input to the dorsal tegmental nucleus (DTN), whose reciprocal connections with the lateral mammillary nucleus (LMN) are thought to form the core HD ring attractor (Blair et al., 1998; Stackman and Taube, 1998; Bassett and Taube, 2001; Sharp et al., 2001b; Taube, 2007; Yoder and Taube, 2014; Graham et al., 2023). This ascending NPH– DTN–LMN pathway has therefore been interpreted as conveying AHV because neurons along it appear AHV-tuned and disrupting NPH, DTN, or LMN destabilizes or abolishes HD tuning (Blair et al., 1998, 1999; Bassett et al., 2007; Butler and Taube, 2015; Butler et al., 2017). Yet these findings show only that the pathway is required for HD stability—not that it carries angular head velocity. This distinction is critical because decades of anatomical, lesion, and electrophysiological work identify NPH not as a dedicated head-motion relay, but as a core oculomotor structure whose neurons encode eye position, eye velocity, and saccade-related signals and contribute to gaze stabilization and control (Lannou et al., 1984; Cullen et al., 1993; Kaufman et al., 2000; Dale and Cullen, 2013, 2015). Thus, two mature literatures assign incompatible computational roles to the same nucleus: the HD field treats NPH as a source of angular head velocity, whereas the oculomotor field identifies it as a premotor integrator for eye movements (Figure 1A).

**Figure 1.**
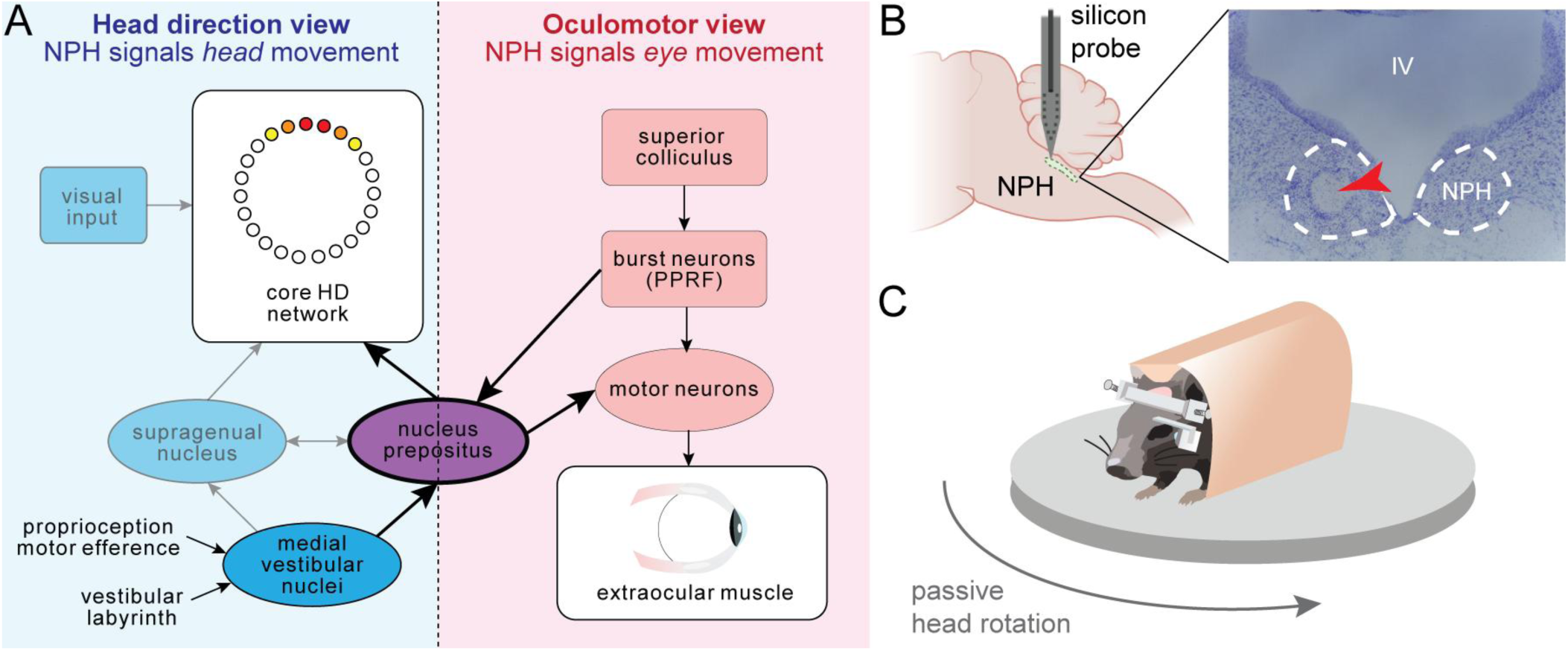
Conceptual background and recording setup. **A)** Blue shading: Canonical models of the rodent head direction (HD) system propose a core HD attractor network, whose activity packet (warm-colored, shaded circles) around a conceptual ring signals the current orientation of the animal’s head. The HD network is thought to receive vestibular information directly from the nucleus prepositus hypoglossi (purple), which moves the activity packet around the ring in response to horizontal rotation. Pink shading: In contrast, the oculomotor literature emphasizes the role of the NPH in horizontal eye movements, particularly through projections to abducens motor neurons. **B)** Setup for **Experiment 1**. 32-channel silicon probes were lowered into the NPH for acute neural recordings during passive head rotation. Electric current was passed through the probe after each session in order to mark the recording location. An example electrolytic lesion mark is shown (red arrow). **C)** Cartoon depicting the behavioral apparatus for the NPH recordings. Mice were implanted with a titanium bar that allowed their head to remain fixed in place while the entire platform was rotated to induce the vestibulo-ocular reflex (VOR). An infrared camera mounted on the platform tracked eye movements. PPRF = paramedian pontine reticular formation.

This conflict has remained unresolved because eye and head movements are tightly coupled. During the vestibulo-ocular reflex (VOR), compensatory eye velocity is approximately equal and opposite to head velocity, so neurons encoding eye movements can appear strongly tuned to AHV when the eyes are not measured. Yet whether NPH explicitly represents head velocity once eye movements are taken into account has never been tested by measuring eye and head movements simultaneously. Most studies identifying AHV-sensitive neurons upstream of the HD system have relied on head kinematics alone, leaving the origin of apparent AHV tuning unresolved (Bassett and Taube, 2001; Sharp et al., 2001b; Graham et al., 2023). More broadly, when correlated behavioral variables are not measured together, neural activity can be assigned the wrong computational meaning. The central unresolved question is therefore what information NPH actually conveys to the HD circuit.

Here we addressed this question by recording from histologically verified NPH neurons while simultaneously measuring eye and head movements in head-fixed mice. Neurons classified as AHV-tuned predominantly encoded eye movements: most showed saccade-related bursts or pauses, and their apparent AHV tuning was largely explained by eye rather than head motion. Optotagging showed that NPH neurons projecting directly to DTN also predominantly encoded eye movements. Empirically constrained population models showed that eye position and gaze direction could be read out directly from NPH activity, whereas recovering a stable head-direction estimate required structured downstream cancellation of its prominent saccade-related signals. Together, these findings overturn the canonical view of NPH as a dedicated AHV relay and reveal eye movements as a major, previously unmeasured component of the signals entering the mammalian HD circuit, thereby changing the computational problem that the HD system must solve to construct a stable representation of heading.

## Methods

This study consists of three main components: two experiments recording from NPH in head-fixed mice, and a computational model. **Experiment 1** tested whether neurons conventionally classified as AHV-sensitive also carried slow-and quick-phase eye-movement signals and used optotagging to identify NPH neurons projecting to the dorsal tegmental nucleus (DTN). **Experiment 2** independently dissociated eye and head motion to determine whether apparent AHV tuning could be explained by eye movements alone. For both experiments, analyses included saccade tuning (peri-event histograms and significance testing), head and eye velocity tuning, and generalized linear modeling. The **computational model** takes the combined data from these experiments to simulate different NPH input connectivity patterns into the head direction compass and tests how angular path integration accuracy is affected.

### Experiment 1: Subjects and timeline

We used male and female mice from Jackson Labs; either wild-type C57BL/6 (“WT”, strain #000664), Ai32(RCL-ChR2(H134R)/EYFP) (“Ai-32”, strain #024109), or Ai27(RCL-hChR2(H134R)/tdT)-D (“Ai27”, strain #012567) mice on a C57BL/6 background. Ai32/Ai27 mice express a channelrhodopsin-2/EYFP fusion protein following exposure to (virally delivered) Cre recombinase. Founders were acquired from Jackson Labs and thereafter bred in-house. Mice were housed in a shared vivarium on *ad libitum* chow and maintained on a 12:12 light/dark cycle. Experiments took place during the light phase. The experimental timeline for each mouse consisted of (1) headbar surgery, (2) following recovery, habituation to the head-fixed setup, (3) craniotomy surgery, and (4) acute recordings (typically 4-6 days). Mice were at least 3 months old at the time of initial surgery and spanned 3–12 months in age at the time of electrophysiological recordings. See Table S1 for breakdown of sex, genotype, and number of recording sessions.

### Experiment 1: Surgical procedures

All procedures were approved by the Dartmouth College Institutional Animal Care and Use Committee. WT and transgenic mice underwent surgery to implant a custom-made titanium headbar for later head fixation. Mice were sedated with 1%-2% isoflurane. After being secured in the ear bars, they were given ketoprofen (analgesia) and Baytril (antibiotic), and dexamethasone for inflammation. After removing the scalp, the headbar was placed horizontally on the skull and cemented in place with C&B Metabond dental cement (Parkell). A small, stainless steel ground screw was screwed into the parietal bone. A small mark was made over bregma and over NPH (-6.2mm posterior) with an ultra fine sharpie pen. A thin layer of transparent Metabond cement was applied to the exposed skull to prevent infection.

Prior to headbar placement, transgenic mice received an intracranial injection into the dorsal tegmental nucleus (DTN) of a retrogradely transported AAV encoding EGFP-Cre under the synapsin promoter (pENN.AAV.hSyn.HI.eGFP-Cre.WPRE.SV40, plasmid #105540-AAVrg, Addgene). Retrograde transport from DTN induced Cre-dependent channelrhodopsin expression in NPH neurons projecting to DTN. A small burr hole was made over the DTN at-5.2 mm posterior to bregma and 0.0 mm mediolateral. Mice were injected with approximately 20 nl of rAAV at a depth of 3.4mm ventral to the skull surface. A removable silicone sealant (Kwik-Cast, World Precision Instruments) was placed over the burr hole. Injection placement in DTN and channelrhodopsin expression in NPH were subsequently confirmed histologically. All mice received 1mL of sterile saline before the end of surgery to maintain hydration. Ketoprofen and Baytril were administered at 24 h and 48 h after surgery. Mice were allowed to recover for one week after headbar surgery before beginning habituation training.

### Experiment 1: Apparatus

The recording apparatus consisted of a custom-built platform with a 3-D printed plastic running wheel, posts (Thorlabs, TR10 & TR6-P5) and clamps (Thorlabs, RA90 & PC2, for securing the headbar in place, and a stereotactic arm (Kopf) for holding the recording probe; Figure 1C). The base of the platform contained a circular array of ball bearings that allowed it to rotate horizontally. The axles of the platform and the running wheel housed rotary encoders (Grayhill 63R256) that were connected to a Teensy 4.0 microcontroller (Sparkfun Electronics, DEV-15583), which in turn was patched into the neural recording system (Digital Lynx SX, Neualynx) as a set of analog signals.

The platform was equipped with a 1.3 megapixel GigE monochrome camera (Basler ace, acaA1300-75gm) for eyetracking. The camera was oriented on the horizontal plane facing the mouse’s left eye. Video was captured via the Cheetah acquisition software (NeuraLynx, Bozeman, MT). The camera had a screw-on longpass infrared (IR) filter (Thorlab, FELH0800) and the mouse eye was illuminated with an IR light source. Recordings were conducted with room lights on.

### Experiment 1: Habituation

Mice were habituated to tolerate head fixation before electrophysiological recordings began. Over the course of several days, the experimenter placed the mouse on their lap and would use their hands to hold the headbar in place for increasing intervals of time as the mice began to tolerate this constraint. Finally, the mice would undergo two sessions in which they were fastened onto the headbar clamps and allowed to move on the running wheel for about 15 minutes.

### Experiment 1: Craniotomy

The day before recording was set to begin, mice were anesthetized and secured in the stereotactic frame. They were given analgesia, antibiotics, and an anti-inflammatory, as before. A small trephine (1.85mm diameter) was used to make a hole in the skull over NPH. The craniotomy was closed with Kwik-Cast and reapplied between recording sessions.

### Experiment 1: Acute recordings with silicon probes

A single 32-channel silicon penetrating probe was used for each recording session (Neuronexus, Ann Arbor, MI), with a tetrode configuration: model *A4×2-tet-5mm-150-200-121*. The probe was connected to a Digital Lynx SX data acquisition system (Neuralynx). Putative action potentials were recorded as threshold crossings of 600 to 6,000 Hz band-pass filtered data sampled at 32 kHz. Signals were referenced locally to the quietest channel.

Once the probe was placed on the stereotactic arm, the probe ground wire was attached to the mouse’s skull screw ground pin. On some sessions, red dye (DiI, Invitrogen) was delicately painted onto the probe shanks with a dental applicator brush (Metabond); (1) either before the recording session, or (2) immediately after a successful recording, in which case the probe was lowered to the same depth. During probe insertion, saline was added periodically to keep the craniotomy moist. During insertion, the experimenter took note of when the ‘tetrodes’ became quiet, indicating the 4^th^ ventricle, and when activity re-appeared, indicating NPH. Recordings were made when one or more well-isolated single units were found below the ventricle.

### Experiment 1: Stimulus and timeline

During a recording session, the platform was periodically rotated manually in a sinusoidal fashion for ∼10 cycles at a time. Periods of rotation were alternated with platform-stationary periods. Efforts were made to capture both slow and fast horizontal rotation speeds. Maximum speeds were at least 100 deg./s, and most sessions had sufficient samples at 150-200 deg./s, allowing comparison of our tuning curves with prior work on the head-direction system in rodents (Graham et al., 2023).

### Experiment 1: Microlesions

After a recording was complete, the headstage and ground wire were carefully removed, and a breakout board (EIB-36 Plating Adaptor, Neuralynx) was attached to the probe. Using a stimulus isolator, current (typically 20 ***μ***A for 15 sec.) was passed through a channel that had a good single unit. Probes were coated with iridium oxide or Z-coat (proprietary formula, Neuronexus) by the manufacturer to increase charge capacity and reduce damage to the probe from electrolytic current.

### Experiment 1: Optotagging

For optotagging sessions, a probe with attached optic fiber was used (Neuronexus, A4×2-tet-6mm-150-200-177-OCM32LP). The fiber was either 100 ***μ***m or 200 ***μ***m in diameter. The light source was a blue diode-pumped solid state (DPSS) laser (Shanghai Laser & Optics Company, BL473T3-100) that was coupled to the probe with a patch cable and plastic mating sleeve (Thorlabs). A light power level of 5-7 mW at the fiber tip was used during the optical stimulation part of the recording protocol. During a recording session, once the probe was in NPH and a candidate neuron was found, a sequence of light pulses was used to detect a visible optically-evoked response. If an optically responsive neuron was found, then recording immediately proceeded and pulse trains were delivered intermittently during platform-stationary periods. Each pulse train consisted of five 1 s long pulses, with a 1 s interval between each pulse end.

Neurons were classified as optotagged when their mean firing rate during the 1-s laser pulse exceeded the 95th percentile of a null distribution generated by circularly shifting the laser-onset sequence relative to the spike train 2,000 times. Classification was based on this shuffle criterion; laser-response latency was quantified separately and was not used as an additional inclusion criterion. Five neurons met the statistical criterion and were histologically verified to lie within NPH. For each optotagged neuron, response latency was calculated as the median latency to the first spike after laser onset and corrected for the independently measured delay between the TTL command and opening of the laser shutter.

### Experiment 1: Histology and recording site localization

After several recording sessions were performed in a given mouse, the mouse was euthanized and perfused with 4% paraformaldehyde (PFA) solution in phosphate buffered saline (PBS). The brain was extracted and placed in PFA overnight then immersed in 30% sucrose in PBS, as a cryoprotectant. After freezing, it was sliced in 40-60***μ***m thick sections on a dry-ice microtome (Leica SM2010R). Half of the sections were stained with thionin to visualize the electrolytic lesions. Alternate sections were mounted onto slides and coverslipped with Vectashield with DAPI (Vector Labs) to visualize the syringe track and any electrode tracks with DiI, as well as viral expression in DTN and NPH (for transgenic mice).

Thionin-stained sections were imaged with a stereomicroscope (Leica M205C) and DAPI-stained slides were imaged with a Keyence BZX-800 fluorescence microscope. The dataset used here (*n* = 104 neurons) is restricted to the subset of recordings in which there was an identifiable electrolytic lesion mark (Figure 1B; see Figure S1 for more examples) or dye-labeled probe track in the NPH. The locations of electrode recording sites were transposed onto mouse atlas images (Paxinos and Franklin, 2001). For recordings in which one electrode site was used to make a lesion, but neuron(s) were recorded on other tetrodes within NPH, the expected location of those recording sites was added with custom MATLAB code, based on the geometry of the probe (for graphic of all recording sites, see Fig. S2).

### Experiment 1: Eye trackin

Eye movements were recorded from an infrared camera situated near the mouse’s left eye at 50Hz (50 sessions) or 33Hz (4 sessions) and Facemap (Syeda et al., 2024) was used to calculate pupil position offline. In order to convert the pupil position time series from pixel space to degrees of visual angle, we took advantage of the fact that AHV and slow phase eye velocity (sEV) are almost perfectly anti-correlated, with studies indicating that sEV amplitude is roughly 80%-100% of AHV amplitude, with a sign change (Hess et al., 1985; Meier and Dieringer, 1993). Based on prior measurements showing that the VOR gain in mice under these conditions is approximately 0.9 (Beraneck et al., 2008), the inverted AHV trace was scaled accordingly and compared to the slow-phase eye velocity (sEV) trace. A calibration factor converting pupil displacement from pixels to degrees was then adjusted iteratively until the amplitudes matched. Once the calibration factor was determined, the EV values were multiplied by this factor to obtain pupil position in degrees. This calibration procedure was used solely to convert pupil displacement from pixels to degrees and did not impose any relationship between neural activity and eye or head kinematics. The relative explanatory power of eye-and head-movement variables was tested independently in Experiment 2 using paradigms that dissociated eye movement from head motion.

The instantaneous eye velocity (EV) signal was calculated from the first derivative of the eye position (EP) signal. Both the eye position and the eye velocity signals were then upsampled to 1 kHz. EV was thresholded to detect both temporal and nasal candidate saccade event times, curated by the experimenter to exclude false-positives and include missed saccade events, and finally re-aligned to saccade onset. Eye movements outside the platform rotation periods (i.e., platform stationary period, AHV=0) were analyzed separately. The evoked eye-movement signals within the platform rotation periods were categorized into two phases: the quick phase component (50 ms before and after each saccade start time) and the slow phase component (excluding quick phases). Slow phase EV and EP signals were filtered using an 11-point median filter and then smoothed using the MATLAB *smoothdata()* function.

### Experiment 1: Neural data preprocessing

Spike waveforms were clustered automatically using KlustaKwik (Harris et al., 2000) and then manually curated using MClust 3.5 (Redish, 2025). Spike trains were imported into MATLAB and analyzed using custom-written algorithms.

### Experiment 1: Neural correlates of quick-phase (saccades)

The eye-movement response of a given NPH neuron was primarily examined by its peri-event time histogram (PETH). Separate temporal and nasal saccade PETHs were constructed by stacking all the saccade event segments for that saccade type together within a window of 200 ms before to 100 ms after saccade onset. This window was large enough to capture baseline firing, any phasic change, and return to baseline. To quantify the degree of firing rate change (either positive or negative), we calculated the absolute value of the difference between the average firing rate in this window and the instantaneous firing rate at each point. The time point with the maximum firing difference (from the average) was considered the peak (or trough) of the phasic response. As a control comparison, we implemented a random circular shuffle of the saccade times over 1000 iterations and calculated the maximum firing rate difference in the same way for each iteration. We compared each neuron’s true phasic response to the distribution of its shuffled phasic responses, and used the 95th percentile as the significance threshold. Thus, a given neuron could have significant quick phase modulation for either saccade type, for both, or neither. The *preferred saccade direction* of a given NPH neuron was defined as the saccade direction with the higher response peak.

*PCA analysis of saccade PETHs:* To further characterize the peri-saccade modulation of firing rates across the population of recorded neurons, we performed principal component analysis (PCA) across the full matrix of preferred and non-preferred saccade PETHs for each cell to obtain a set of principal components (PCs). We projected each cell’s preferred and non-preferred saccade PETHs onto the first principal component to obtain the coefficient (loading) on the first PC, and plotted these coefficients against each other in Figure S4.

### Experiment 1: angular head velocity (AHV) tuning analysis

Angular head velocity was calculated by taking the derivative over time of the orientation of the recording platform. In order to directly compare our recordings with previous work, we used data analysis steps and the classification criteria from Graham et al., 2023 to categorize cells as AHV-sensitive. The mean velocity and firing rate for each cell was binned in 6°/s bins, with all data above 204°/s being excluded. A session needed to have velocities of at least 150°/s to be analyzed. Individual bins with less than 0.5 s of data were excluded. Cells were classified as AHV-sensitive if they met three criteria: (1) The best-fit line for the two halves (CW and CCW) of the tuning curve contained a Pearson correlation ≥ 0.5, (2) the absolute value of the slope of the best-fit line was ≥ 0.025 spikes/°/s, and (3) each cell needed to pass a shuffle test. For the shuffle test, the spike train for each neuron was randomly shifted (min shift = 5 s) relative to the position tracking data 1000 times. For each shuffled spike train, a new AHV tuning curve was generated and the best-fit line slope calculated for the two halves of the tuning curve. To pass the shuffle test, the true slope on either the CW or the CCW side needed to rank above the 95th percentile of the shuffled values.

*Categories of AHV-selective cells*: AHV cells were divided into categories that described their tuning curve profiles. We calculated a “turn bias index” using the same formula as Graham et al., 2023:

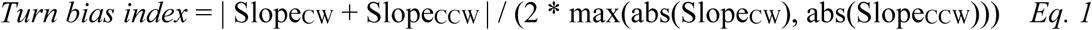

where Slope_CW_ is the slope of the binned, average tuning curve for the CW direction. Slope_CCW_ is the slope of the binned, average tuning curve for the CCW direction. According to this formula, cells that are perfectly asymmetric will have values near 1. Cells with values near zero (≤ 0.3) were defined as “symmetric.” Cells with values ≥ +0.7 were defined as “asymmetric.” Cells that had values between +0.3 and +0.7 were defined as “rectified,” meaning that half of the tuning curve was sloped, and half of the tuning curve was flat.

### Experiment 1: Neural correlates of slow-phase analysis

To further quantify the eye-movement responses of individual NPH neurons, we fit an eye-kinematic generalized linear model based on Sylvestre and Cullen (1999):

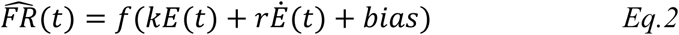

where *FR*^(*t*) is the neuron’s estimated firing rate; *E*(*t*) is the eye position; *Ė*(*t*) is the eye velocity; *bias* is the resting discharge; *k* is the eye position sensitivity, and *r* is the eye velocity sensitivity. The model assumed a normal response distribution with an identity link function. For each model fit, we determined the variance accounted for (VAF) as

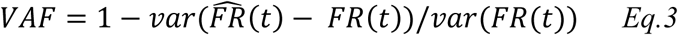

where *FR*(*t*) is the measured firing rate. A *VAF* of 1 indicates a perfect prediction, whereas a *VAF* of 0 indicates that the model performs no better than predicting with the neuron’s mean firing rate. Periods when the platform was stationary (AHV = 0) were discarded. For modeling the slow phase, all data within 50 ms of a saccade were removed. For modeling quick phase, only data within 50 ms of a saccade were used. The model fitting was repeated 2000 times with random sample bootstrapping, and the averaged eye position sensitivity, eye velocity sensitivity, resting discharge, and VAF were reported. 95% bootstrap confidence interval threshold for *k* and *r* were used to classify neurons as significantly sensitive to eye position, eye velocity, both variables, or neither.

### Experiment 2: Subjects and timeline

Five male C57BL/6J (The Jackson Laboratory) adult mice were included in this study. All procedures were conducted under approved animal protocols at Johns Hopkins University.

### Experiment 2: Head-post implantation and craniotomy

Comparable to Experiment 1; briefly, mice were anesthetized with isoflurane 2% in oxygen. A 1.5 mm-diameter craniotomy was performed. A custom-built head-post was cemented on the skull. Bupivacaine (0.5%) was given before and after surgery. Care was provided to avoid hypothermia and dehydration.

### Experiment 2: Electrophysiological recording

Neural activity of mouse NPH neurons was collected using either tungsten electrodes (8–10 MΩ impedance, FHC) or Neuropixels probes (IMEC). To locate the nucleus prepositus, we first located the abducens nucleus based on the characteristic discharge of its neurons to ipsilateral eye position/velocity during visually evoked slow phase movements and bursts of action potentials for ipsilaterally directed saccades (Stahl and Thumser, 2012). We then mapped the nucleus’ anterior–posterior and medial–lateral boundaries, verifying that its center comprised ∼200–250 μm of this characteristic activity along a single recording track. All neurons in this study were recorded at a location ∼200 μm posterior to the center of the abducens nucleus, in the anterior NPH.

Tungsten electrodes were used with a custom-built microdrive. Single-unit activity from tungsten electrodes was recorded through a Plexon system at 20 kHz for offline analysis. For acute Neuropixels recordings, an implant assembly was developed that allowed flexible probe positioning and retraction after experiments. The probe depth from skull surface was ∼4.5 mm.

Neuropixels data were collected using 1.0 single-shank probes (Jun et al., 2017) and SpikeGLX (Bill Karsh, https://github.com/billkarsh/SpikeGLX) at a sampling rate of 30 kHz.

### Experiment 2: Recording setup

Eye and head movements were recorded simultaneously with neural activity. The mouse was placed in a body tube and head-fixed. It was then positioned on a rotating turntable surrounded by a drum with vertical black-and-white stripes with a spatial frequency of 0.1 cycles per degree (Fig. 5). Head movements (whole-body rotation) were measured using a gyroscope on the turntable (MPU-9250, SparkFun Electronics). Because the head was fixed relative to the turntable, head motion refers here to horizontal whole-body/platform rotation in yaw. Eye movements were measured using either (1) a magnetic eye tracking system (Payne and Raymond, 2017) or (2) a mini camera system (Stahl, 2004) at a sampling rate of 50 Hz.

### Experiment 2: Behavioral paradigms

Responses of nucleus prepositus neurons were examined using three paradigms designed to either couple or dissociate eye and head movements (Fig. 5A). To isolate eye-position signals from head motion, we first used the static eye position (SEP) paradigm (Beraneck and Cullen, 2007), in which the platform was rotated through a series of discrete steps and then held stationary. Following each step, the eyes stabilized at different orbital eccentricities spanning an ∼20° range. To estimate eye-velocity sensitivity in the absence of head movement, we next measured neural responses during the optokinetic reflex (OKR), evoked by rotating the striped drum while the turntable remained stationary. OKR stimuli consisted of sinusoidal rotations at 0.5 Hz (40°/s peak velocity) and 0.2 Hz (20°/s peak velocity). Finally, vestibulo-ocular reflex (VOR) responses were measured by rotating the turntable while the visual surround remained stationary, thereby evoking coupled head and compensatory eye movements. VOR stimuli consisted of sinusoidal rotations at 0.5 Hz (40°/s peak velocity) and 0.2 Hz (12°/s peak velocity).

### Experiment 2: Data analysis

Raw spike signals were imported into MATLAB and analyzed using custom-written algorithms. Neuropixels data were first sorted using Kilosort (Pachitariu et al., 2016) and curated with Phy (https://github.com/kwikteam/phy). As in Experiment 1, eye position and velocity sensitivities were computed using a generalized linear firing-rate estimation model and model performance was quantified using VAF.

Models were first fit to static eye-position (SEP) and optokinetic reflex (OKR) responses, conditions in which eye position or eye velocity varied in the absence of head motion. The coefficients estimated from these eye-only conditions were then applied directly without refitting to predict each neuron’s firing rate during the vestibulo-ocular reflex (VOR) condition, when matched eye movements occurred together with passive head rotation. This cross-condition analysis tested whether eye-movement sensitivities estimated in the absence of head motion were sufficient to explain neural responses during head motion.

For the VOR condition, residual firing rate from the eye kinematic model prediction was calculated as

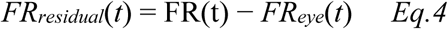

where *FR^_eye_*(*t*) was generated using coefficients estimated from SEP and OKR. Residual firing rate was then fitted with head velocity to test whether head motion explained additional neural variance beyond the eye-movement model using a head-kinematic generalized linear model:

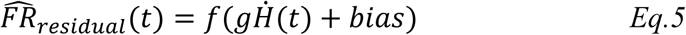

where *FR*^*_residual_*(*t*) is the neuron’s estimated firing rate residual; *Ḣ*(*t*) is the head velocity; *bias* is the offset term; *g* is the head velocity sensitivity, For each model coefficient computed during SEP, OKR, and VOR conditions, 95% bootstrap confidence intervals were generated (Dale and Cullen, 2013). Specifically, 2,000 datasets were created by resampling with replacement. Coefficients whose 95% confidence intervals did not overlap zero were considered significant. Eye-position and eye-velocity sensitivities were also compared between matched OKR and VOR conditions to determine whether introducing head rotation altered the estimated oculomotor sensitivities.

### Model: Simulation of pseudo-simultaneous NPH populations

The overall objective of the modeling component was to determine what signals could be read out from the empirically observed tuning of NPH neurons. To this end, eye-position and eye-velocity sensitivities, and bias coefficients during slow phase and quick phase estimated from the recorded NPH neurons were used to generate model-based firing-rate responses during simulated VOR, OKR, and spontaneous-saccade conditions. NPH neurons were identified as “same” or “opposite” class according to whether their preferred quick-phase and slow-phase response directions were aligned or opposed, respectively. Pseudo-populations with total neuron number N of 10, 20, 30, 40, and 80 were generated by sampling recorded neurons with replacement while preserving a series of proportions from 100%:0% to 0%:100% between the “same” and “opposite” classes. For each total neuron population number N and each “same” versus “opposite” class ratio, the pseudo-population were generated with 50 repetitions with random seed values.

For each simulated neuron in a pseudo-population, the model-predicted firing rate was converted into a spike train using a nonhomogeneous gamma process. A good approximation of nonhomogeneous gamma process was realized by applying a thinning procedure onto a high-rate homogeneous gamma process (Lewis & Shedler, 1979). At any time *t*, the interval to the next spike candidate *δt* was sampled from a high-rate homogeneous gamma process using the MATLAB *gamrnd*() function and added to the time *t*:

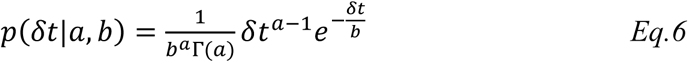

Where *a* is the shape parameter that determines interspike-interval regularity, and *b* is the scale parameter that determines the mean interspike interval. We chose the shape parameter *a* = 2 for slow-phase spike simulation and *a* = 100 for quick-phase spike simulation to capture the different firing patterns associated with the relatively smooth modulation during slow-phase eye movements and the abrupt burst–pause responses during quick phases. The scale parameter was set so that the homogeneous gamma process maintained a constant rate corresponding to the maximum bound of the NPH firing rate. In addition, a minimal 1-ms refractory period was set to prevent biologically impossible spike burst activities. After spike candidate was generated from the high-rate homogenous gamma process, thinning procedure was applied to accept or reject this spike by independently checking whether a random variable sampled from uniform distribution *u*∼*U*(0,1) falls within a spike firing threshold, which was calculated as the model-predicted firing rate at time *t* divided by the maximum bound firing rate:

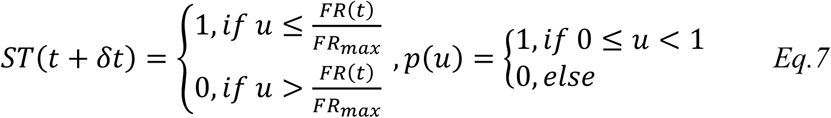

Choosing gamma process allows us to better represent NPH neuronal spike train properties such as regularity and refractory period, comparing the fixed exponential shape from Poisson process.

The resulting pseudo-simultaneous spike trains preserved the empirically measured relationships among slow-phase, quick-phase, and bias terms while allowing population size and response-class composition to be manipulated systematically.

### Model

Population decoding of orientation variables. The pseudo-population of simulated NPH spike trains were provided as inputs to a network model that decoded head direction, eye position, or gaze direction. The corresponding excitatory postsynaptic current (EPSC) was calculated by convolving each spike train with a normalized alpha-function kernel:

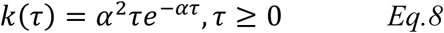

The EPSCs from multiple NPH neurons were combined as a weighted sum with a constant offset, with weights estimated using non-negative linear regression, to derive the network driving signal (see Eq. 3 below). The downstream ring-attractor dynamics were approximated as temporal integration of this population driving signal to generate the decoded orientation estimate.

The optimized weight mapping of the NPH neuron population was determined by minimizing the squared estimation error as follows:

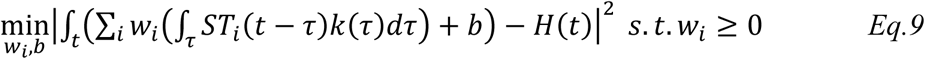

where *ST_i_*(*t*) is the binary spike train of the *i*th NPH neuron at time *t*, *k*(*τ*) is a normalized alpha-function kernel, with a 15 ms time constant. *H*(*t*) is the target orientation variable, *w_i_*is the weight assigned to the *i*th NPH neuron, and *b* is the constant offset. Thus, *b* served as an intercept term. All neuronal weights were constrained to be non-negative, consistent with prior models (Song and Wang, 2005).

Decoder weights were optimized for each pseudo-population. For each evaluation, one session was held out as a test set and the remaining sessions were used for training. Signals from the training sessions were segmented into 1-s epochs and pooled to estimate decoder weights by minimizing the squared estimation error. Model performance was then evaluated on the held-out session using these fixed weights, ensuring that no test data were used during model fitting. Decoding error was quantified as the sample averaged square root of the squared difference between the integrated population drive and the target orientation variable. To test cross-context generalization, decoder weights optimized during simulated VOR were held fixed and applied without refitting to simulated OKR and spontaneous-saccade activity.

## Statistical analysis

Statistical analyses were performed using MATLAB. Data are reported as mean ± SEM unless otherwise indicated.

Shuffle tests (random circular shifts of spike times or laser onsets relative to the behavioral or stimulus signal) were used to assess significance for three analyses: (1) peri-saccadic firing-rate modulation in Experiment 1 (1000 iterations per cell; 95th percentile threshold on the maximum firing-rate difference); (2) AHV-tuning classification in Experiment 1 (1000 iterations per cell; 95th percentile threshold on best-fit slope of the CW or CCW half of the tuning curve); and (3) laser-evoked firing during optotagging (2000 iterations per cell; 95th percentile threshold on mean firing rate during the 1 s laser window).

Bootstrap resampling with replacement (2000 iterations) was used to compute 95% confidence intervals on eye-position and eye-velocity sensitivity coefficients from the generalized linear model fits in Experiments 1 and 2, and to estimate uncertainty for VAF. Wilcoxon signed-rank tests were used for paired comparisons of resting discharge, eye-position sensitivity, and eye-velocity sensitivity between matched OKR and VOR conditions in Experiment 2. The Wilcoxon signed-rank test was used for paired within-cell comparisons (e.g., preferred-direction burst amplitude during VOR saccades vs. spontaneous saccades, Fig. S5).

A 2-way ANOVA (factors: behavioral context, population size) was used for comparisons of decoding error and generalization ability between head direction and eye position decoding (Fig. 8).

## Results

### Experiment 1: NPH neurons exhibit apparent angular head velocity tuning

To directly compare our data with prior studies of the head direction system, we first analyzed NPH activity using the conventional framework of angular head velocity (AHV) tuning. In **Experiment 1**, mice were passively rotated through a range of angular velocities, thereby evoking the vestibulo-ocular reflex (VOR), while neurons were recorded from NPH. To facilitate comparison with prior HD studies, we plotted each neuron’s tuning curve as a function of AHV (Fig. 2A). Firing rates showed strong apparent modulation with AHV. Using the classification criteria of Graham et al., (2023), we identified neurons that would conventionally be classified as AHV-responsive (see *Methods*). A large majority of neurons met these criteria (84/104, 81%; 55% ipsiversive, 45% contraversive relative to the recording hemisphere). We further classified the shapes of their tuning curves: 30 (28.8%) asymmetric, 49 (47.1%) rectifying, 5 (4.8%) symmetric, and 0 were inverted.

**Figure 2.**
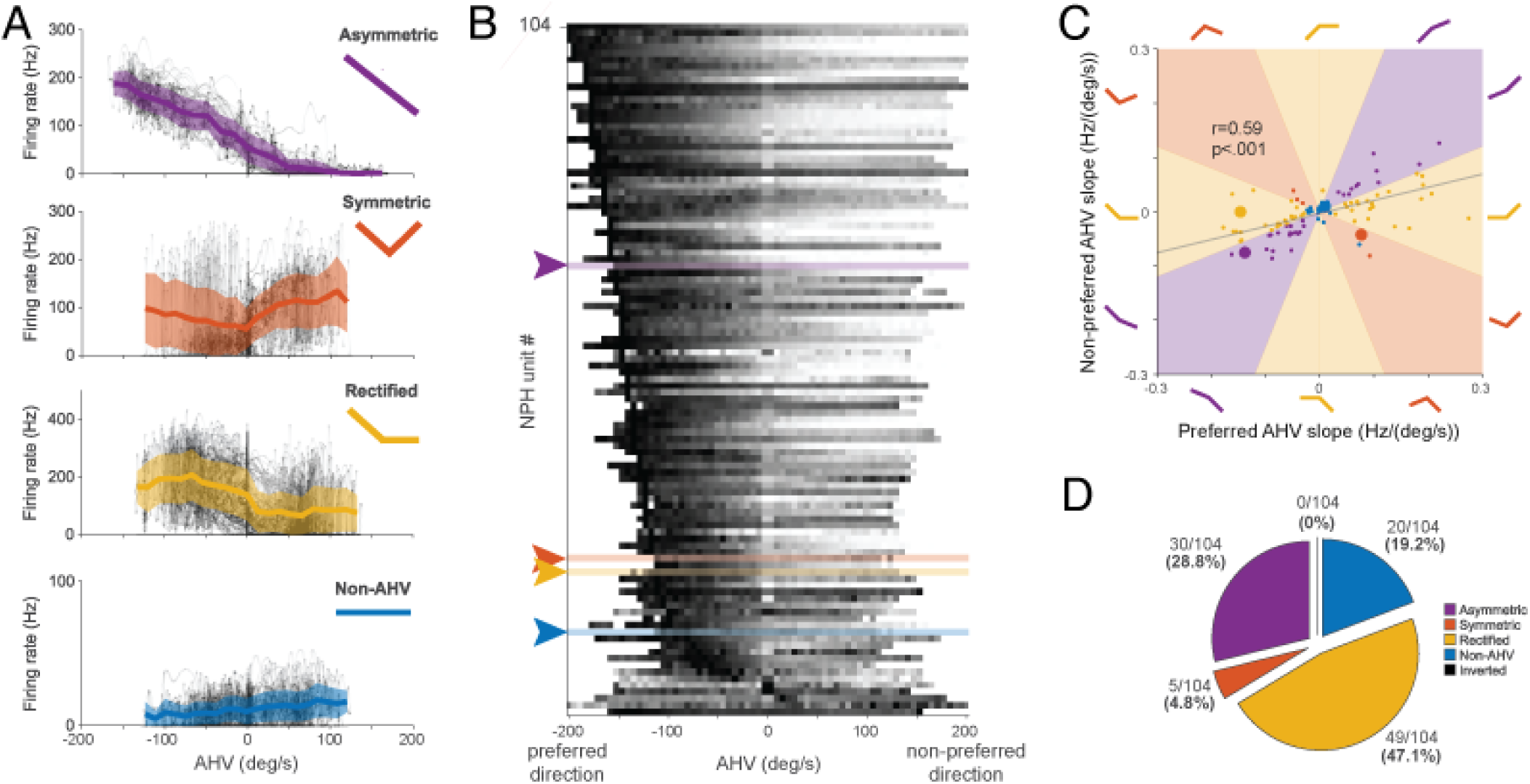
NPH neurons exhibit canonical AHV-like tuning profiles. **A)** Example AHV tuning curves. Gray dots show individual data points. Solid lines represent the mean ±SEM. First row: “asymmetric” AHV cell with a significant linear fit for CW and CCW directions. Second row: V-shaped “symmetric” cell that increases firing rate during turns in both directions. Third row: “Rectified” cell with significant slope for one direction only. Fourth row: “Non-AHV” cell. There were no “inverted” cells with an upside-down V-shape tuning curve. **B)** Stacked heatmap showing normalized AHV tuning curves for all NPH neurons (one neuron per row), and arranged according to the bin with maximum firing rate. For visualization, each tuning curve is plotted with the preferred direction *on the left side*. Example neurons from panel A are indicated by arrows and corresponding colors. **C)** Scatterplot of preferred-direction (abscissa) and non-preferred-direction (ordinate) slopes, one data point per neuron. Large, colored circles correspond to the example cells shown in panel A. **D)** Breakdown of AHV response categories. Colors correspond to the example neurons shown in panel A. Neurons were categorized as AHV-responsive if they satisfied criteria for correlation, slope, and a shuffle test.

Example cells from each category are shown in Figure 2A. The preponderance of asymmetric and rectifying responses was evident in the population of tuning curves (Fig. 2B) and in the overall correlation between the slopes of the preferred and non-preferred halves of their tuning curves (Fig. 2C). The asymmetry is also evident in the grand-average AHV tuning curve across all neurons (Fig. S3A). Importantly, these rectified and asymmetric response profiles resemble the velocity signals proposed to update the HD ring attractor in canonical HD models. However, because compensatory eye movements are tightly coupled to head rotation during the VOR, apparent AHV tuning alone cannot distinguish whether these responses reflect head motion, eye motion, or both.

### NPH neurons exhibit bidirectional saccade-related modulation

In order to examine the relationship between eye movements and NPH activity, we divided the data during VOR into quick-phase (saccade) and slow-phase eye movement epochs (an example trace is shown in Fig. 3A; note the anticorrelation between head and eye rotation, quantified in Fig. S6). Figure 3B shows raster plots from an example NPH neuron, along with eye position, aligned to preferred (top) or non-preferred (bottom) saccade onset times. This neuron exhibited robust bidirectional peri-saccadic modulation: a burst of spikes before saccades in one direction, and a transient decrease in spiking before saccades in the opposite direction. Overall, 82/104 (79%) of NPH neurons showed significant saccade-related modulation in at least one direction (see *Methods* for significance criteria). Figure 3C shows peri-event time histograms (PETHs) for all NPH neurons aligned to preferred and non-preferred saccades; bidirectional firing rate modulation across the population was clearly visible around saccade onset and was further confirmed by the anti-correlation of principal component loadings for the preferred and non-preferred directions (Fig. S4). Notably, of the 84 neurons classified as AHV-responsive in the previous section, 68/84 (81%) were significantly modulated for saccades in at least one direction. Thus, consistent with the oculomotor literature, saccade-related firing rate changes were common not only across the NPH population as a whole, but also among neurons conventionally classified as AHV-responsive.

**Figure 3.**
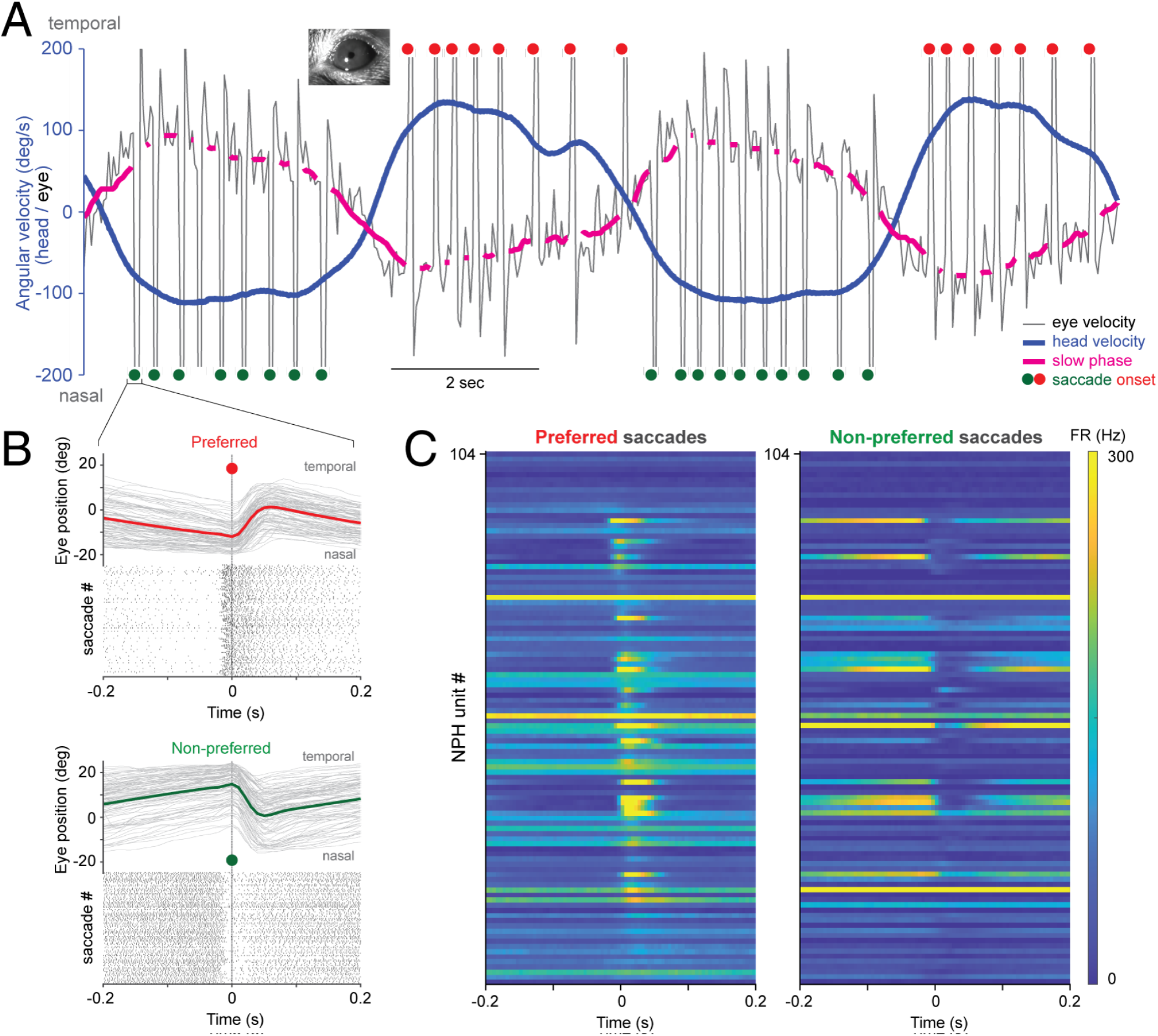
NPH neurons show bidirectional peri-saccadic burst/pause activity. **A)** Example eye velocity trace in black illustrating the two-beat pattern of vestibular nystagmus. A slow-phase component (magenta line) alternates with fast, high-amplitude saccades in the opposite direction, indicated by red (preferred direction for the neuron in panel B) or dark green (non-preferred) circles. Note the anticorrelated relationship between head velocity (blue) and slow phase eye velocity (magenta). Inset: camera view of the mouse’s eye for eye tracking. **B)** Eye position traces during saccades (bold lines indicate trial averages), together with spike rasters from a single neuron aligned to preferred-direction (red) and non-preferred-direction (green) saccades. **C)** Stacked peri-saccade time histograms. Each row displays the firing rate of one NPH neuron for preferred and non-preferred saccade types. Within each panel, neurons are arranged according to the bin with maximum firing rate. Warmer colors indicate higher firing rate. Note the burst of activity preceding and during preferred-direction saccades, and transient suppression of firing preceding and during non-preferred saccades.

Importantly, this peri-saccadic activity was not simply an indirect consequence of vestibular input during the VOR. In the subset of neurons with ≥10 spontaneous saccades per direction during platform-stationary periods (*n* = 28 neurons), peri-saccadic firing rate in the preferred direction was significantly elevated above baseline (median +25.0 Hz; Wilcoxon signed-rank, p = 4.2 × 10⁻⁶; Fig. S5).

### Relationship between quick-phase and slow-phase tuning in NPH neurons

To further characterize the eye-movement correlates of NPH neurons, we next examined their slow-phase tuning. Given that AHV and slow-phase eye movements are anticorrelated during the VOR (Fig. S6), it is not surprising that 79/84 (94%) of AHV-responsive neurons had significant slow phase tuning (Figure 4A), distributed roughly equally between the two slow-phase directions (CCW: 44/79, 56%; CW: 35/79, 44%). The majority (63/79, 80%) of slow-phase sensitive neurons also exhibited significant quick-phase sensitivity.

**Figure 4.**
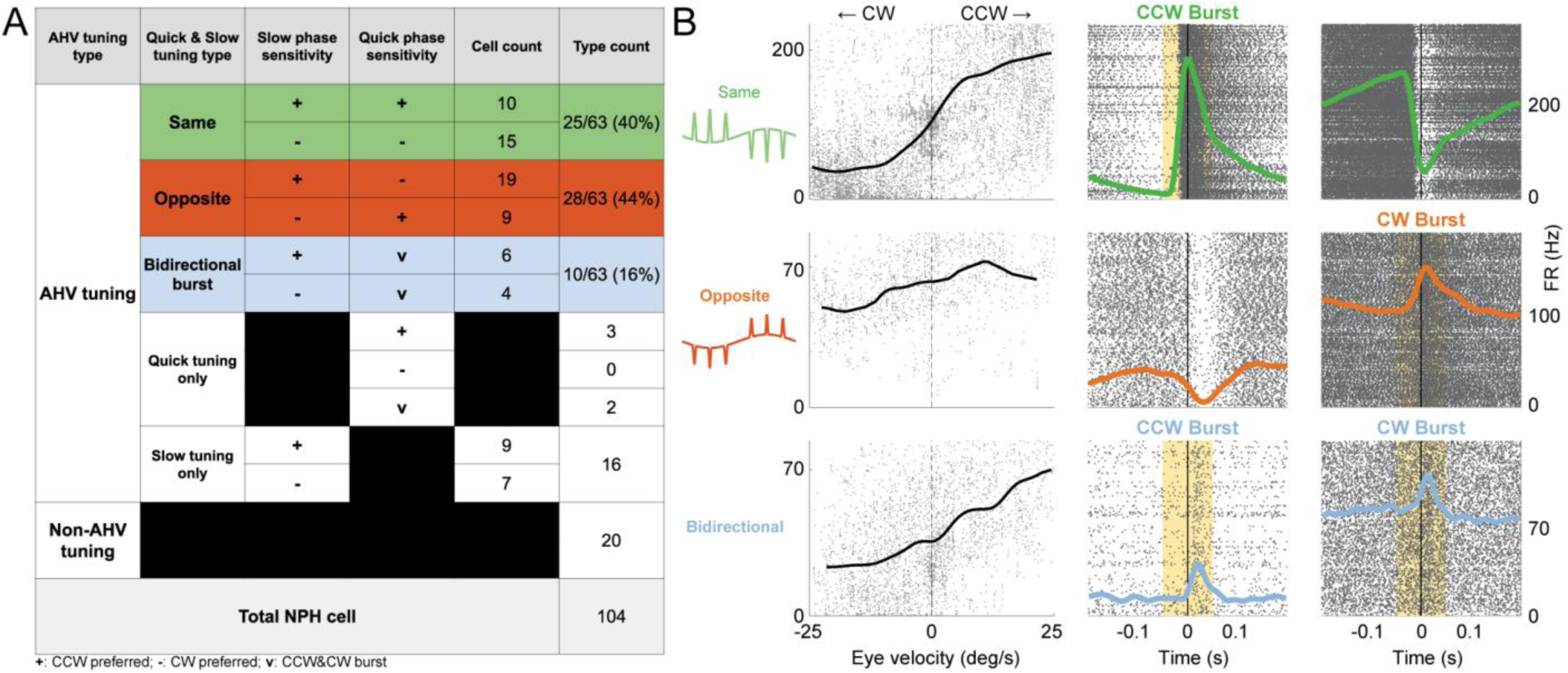
Most NPH AHV-tuned neurons show eye-movement sensitivity. **A)** Table showing the overlap of AHV tuning and eye movement tuning. Among 84 AHV neurons, 75% show both slow and quick phase tuning. Of these, 40% show the *same* preferred tuning direction for slow and quick phase (green); 44% show *opposite* preferred tuning (orange); 16% show *bidirectional burst* quick phase tuning. **B)** Eye velocity tuning curves (**left column**) and peri-saccade raster plots (**middle column:** saccades in CCW direction; **right column**: saccades in the CW direction) for the three different AHV/slow phase/quick phase neuron types. **(top row)** Example *same* type neuron, which prefers CCW slow phase and CCW quick phase. **(middle row)** Example *opposite* type neuron, which prefers CCW slow phase, but CW quick phase. **(bottom row)** Example *bidirectional burst* type neuron, which prefers CCW slow phase and bursts for each direction of quick phase. “+” symbol: CCW-preferring; “-” symbol: CW-preferring; “v” symbol: bidirectional burst.

**Figure 5.**
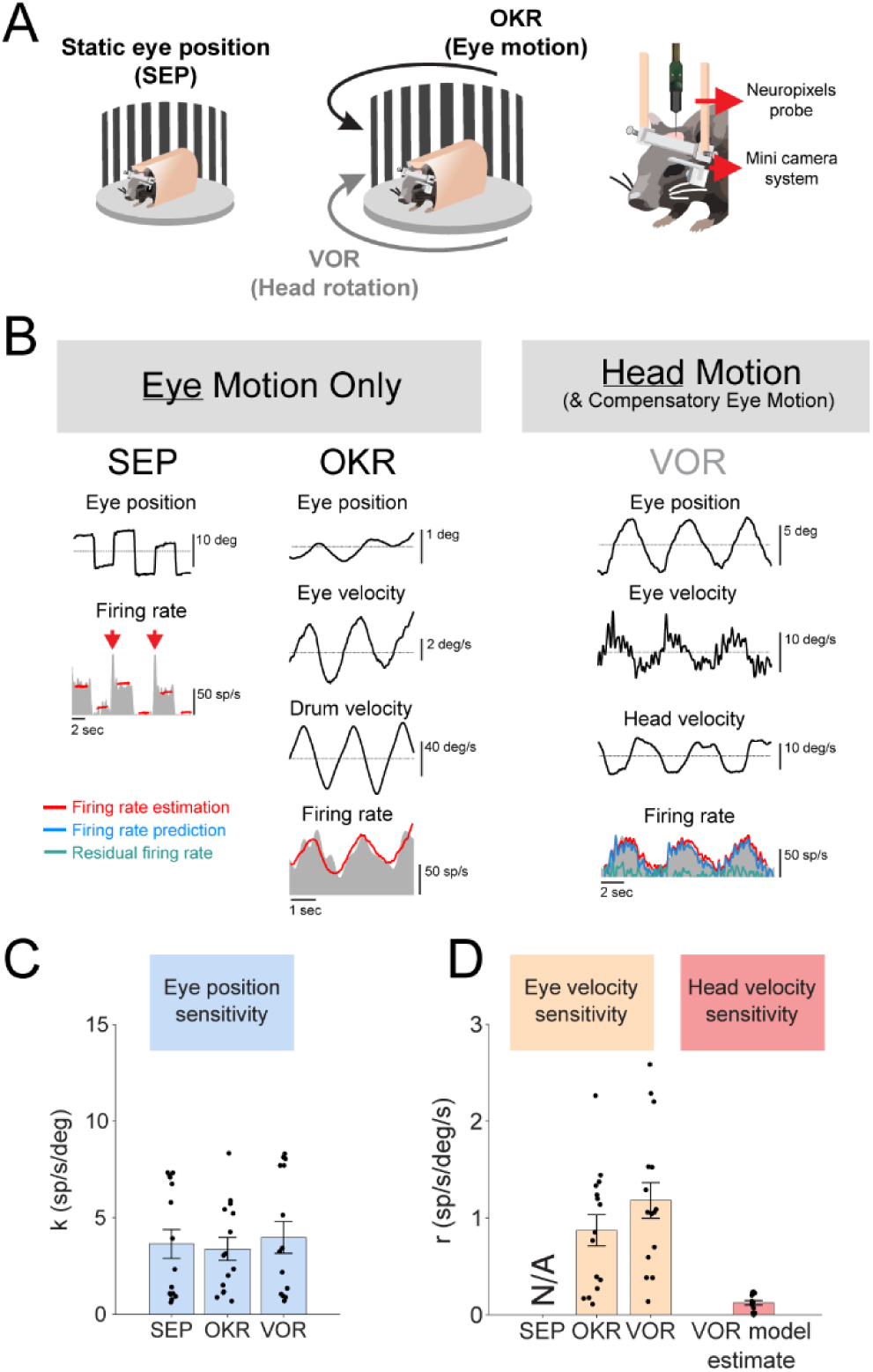
NPH neurons are sensitive to eye position and eye velocity, but not head velocity. **A)** Behavioral apparatus for experimentally isolating eye motion from head motion. Head-fixed mice were seated on a turntable, surrounded by a striped visual drum. Recordings were made from NPH with Neuropixels 1.0 probes. **B)** Schematic showing representative data from the three behavioral paradigms. **(Left)** Static Eye Position (SEP) – the platform was moved in discrete steps to drive eccentric eye positions. **(Middle)** Optokinetic Reflex (OKR) – the drum was rotated back and forth to evoke compensatory eye movements in the absence of head motion. **(Right)** Vestibulo-ocular reflex (VOR) – the turntable was rotated back and forth to evoke passive head motion and compensatory eye movements. Traces to the right of each panel show eye position, eye velocity, drum or head velocity, and firing rate from an example NPH neuron. SEP shows only eye position and firing rate; eye and drum velocity are zero during static eye-position trials. A general linear model was used to calculate the predicted firing rate and the residual firing rate. VOR model predictions were generated exclusively from parameters fit under SEP and OKR conditions (i.e., in the absence of head motion). **C)** Bar graphs showing the firing rate sensitivity in each behavioral condition to eye position (blue). **D)** Bar graphs showing firing rate sensitivity to eye velocity (beige) and head velocity (red). NPH neurons remained strongly sensitive to eye-movement variables even in the absence of head motion, whereas head velocity contributed little additional explanatory power during VOR conditions.

Altogether, 63/84 (75%) of AHV-tuned neurons showed both significant slow-and quick-phase tuning. Among these neurons, the preferred directions of quick-and slow-phase tuning were either aligned (“*same*”, 25/63, 40%; consistent with extraocular motor neuron responses) or opposed (“*opposite*”, 28/63, 44%), or characterized by bidirectional quick-phase bursts irrespective of slow-phase direction (“bidirectional burst”, 10/63, 16%). Figure 4B shows example *same* and *opposite* cells. Thus, neurons conventionally interpreted as encoding AHV instead robustly encoded both slow-and quick-phase eye movements, often in diverse and opposing combinations across the population. Additional analyses further demonstrated that heterogeneous oculomotor coding was a dominant organizing feature of NPH activity (Fig. S7)

### Experiment 2: Apparent slow-phase “angular head velocity tuning” is best explained by eye movements

While the above results unambiguously establish that NPH neurons encode eye movements, the strong anticorrelation between head and eye movements during the VOR leaves unresolved whether head or eye movements are the primary driver of slow-timescale NPH activity during head turns. To directly dissociate these possibilities, we used the behavioral apparatus shown in Fig. 5A, in which mice were head-fixed on a movable turntable surrounded by a striped visual drum (vertical black and white stripes, 5° visual angle width). Eye movements were evoked under three conditions: (1) static eye position (SEP), in which the platform was stepped and then held stationary, producing eccentric eye positions in the absence of ongoing head motion (Fig. 5A, left); (2) the optokinetic reflex (OKR), in which the drum rotated to evoke slow-phase eye movements in the absence of head movement (Fig. 5A, middle, black arrow); and (3) the vestibulo-ocular reflex (VOR), in which the turntable rotated while the drum remained stationary, evoking matched eye movements accompanied by passive head rotation (Fig. 5A, middle, gray arrow). Rotation velocities in OKR and VOR paradigms were adjusted so that the peak velocities of evoked slow-phase eye movements were matched at approximately ±12°/s at 0.2 Hz. We recorded *n* = 15 NPH neurons in these matched OKR and VOR conditions.

We first characterized the firing rates of single NPH neurons while mice generated eye movements in the absence of head motion (i.e., SEP and OKR conditions) and fit a dynamic eye-movement model to the activity of each neuron. A model based solely on eye position and velocity (see *Methods*) accurately described NPH neuron firing rates during SEP and OKR, with a mean population variance accounted for (VAF) of 0.51 ± 0.07 for SEP and VAF=0.58 ± 0.05 for OKR conditions. To directly test whether NPH neurons encode head motion independently of eye movements, we next asked whether the same eye movement-based model, fit entirely in the absence of head motion, could predict neural activity during VOR without refitting. Strikingly, introducing passive head rotation did not improve or alter the model’s predictive power: eye-movement coefficients estimated in the absence of head motion predicted NPH firing during VOR with the same accuracy as during OKR (VOR VAF = 0.58 ± 0.03; OKR VAF = 0.58 ± 0.05). Thus, eye-movement variables alone were sufficient to account for NPH activity during VOR. Consistent with this interpretation, the estimated firing rate bias (i.e. resting discharge rate) did not differ significantly between matched OKR and VOR conditions (OKR vs VOR, 54.5 ± 7.4 vs 58.7 ± 7.3 (spike/s), p = 0.10). Likewise, introducing passive head rotation did not significantly alter either eye-position or eye-velocity sensitivities. Eye position sensitivities were similar between OKR and VOR conditions (Fig. 5C, blue; OKR vs VOR, 3.4 ± 0.6 vs 4.0 ± 0.8 (spike/s)/(deg), p=0.13), as were eye velocity sensitivities (Fig. 5D; OKR vs VOR, 0.9 ± 0.2 vs 1.2 ± 0.2 (spike/s)/(deg/s), p=0.22).

Finally, we tested whether the residual firing activity remaining after subtracting the response predicted by the eye-movement model contained additional head-motion information during the VOR paradigm. Residual firing rates from both OKR and VOR conditions showed negligible modulation to head velocity, with a mean population VAF of only 0.03 ± 0.01. Thus, head velocity provided minimal additional explanatory power beyond the eye-based model. Importantly, head velocity sensitivities during the VOR were an order of magnitude smaller than eye-velocity sensitivities (Fig. 5D, red; VOR head velocity sensitivity=0.12 ± 0.02 (spike/s)/(deg/s)). Together, these results indicate that NPH neurons predominantly encode eye position and eye velocity, whereas head velocity provides little additional explanatory power. Apparent “AHV tuning” in NPH can therefore arise from oculomotor signals that generalize across behavioral contexts, including during head motion.

### DTN-projecting NPH neurons carry eye-movement signals

The above results establish that eye-movement coding is both prevalent across the NPH population (Experiment 1) and quantitatively sufficient to explain NPH firing during head motion (Experiment 2). However, these findings alone do not exclude the possibility that a specialized subpopulation of NPH neurons projecting to the core HD circuit selectively encodes AHV independent of eye movements. Because NPH projects not only to the DTN-LMN pathway, considered the core of the head direction attractor, but also to multiple other targets (McCrea and Horn, 2006; Mehlman et al., 2021), we next asked whether eye-movement signals were specifically encoded by DTN-projecting NPH neurons. To identify DTN-projecting neurons, we used a retrograde viral strategy in which a retrograde AAV carrying Cre recombinase was injected into the DTN of Ai32 mice, resulting in Cre-dependent channelrhodopsin and EGFP expression in projecting neurons (Fig. 6A,B). Consistent with known ascending anatomical projections to DTN, this approach produced robust viral expression in NPH (Fig. 6C) as well in the supragenual nucleus (SGN; not shown). We then repeated the VOR recordings from Experiment 1 using recording probes equipped with an optical fiber. NPH neurons showing significant increases in firing during 1-s blue-light pulses were classified as optotagged DTN-projecting neurons (permutation test; n = 5), and recording locations were confirmed with electrical microlesions and fluorescence microscopy (Fig. 6C,D).

**Figure 6:**
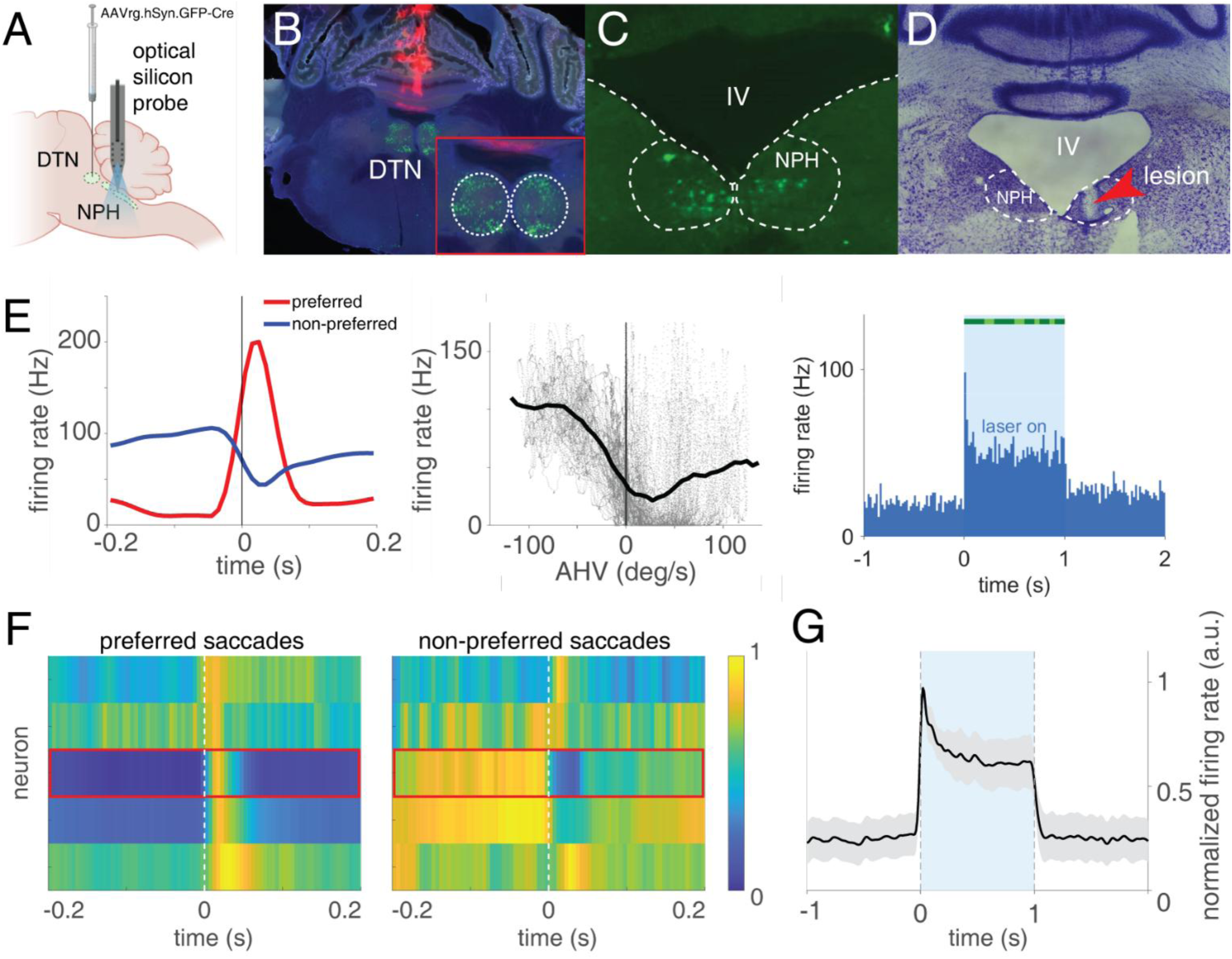
DTN-projecting NPH neurons encode eye movements. **A)** Schematic illustrating the locations of viral injection into DTN and subsequent optoprobe recordings in NPH. Panels B–E present data from a single recording. **B)** Brain slice mounted with DAPI medium (blue) showing DiI-labeled injection track (red) and transfected cell bodies of DTN neurons expressing EGFP. Inset shows DTN at higher magnification. **C)** More caudal brain section from the same mouse showing retrogradely-labeled neuron somata (EGFP) in anterior NPH. **D)** Adjacent thionin-stained brain section showing an electrolytic lesion in the right NPH. **E) Left:** Peri-saccade time histogram (PETH) for the NPH neuron from panel D). **Middle:** AHV tuning curve for the same cell. **Right:** Laser-evoked firing-rate response confirming optotagging. The green bar above the laser-on period indicates time bins with firing rates significantly exceeding a circular-shift shuffle distribution (dark green: p < 0.001; light green: p < 0.05). **F)** Saccade-aligned PETHs for the 5 optotagged cells (all of which exhibited significant quick-phase tuning), shown as a heatmap. **Left:** preferred-direction saccades. **Right:** non-preferred-direction saccades. The neuron from panels D–E is indicated with a red box. **G)** Population-average PETH showing the average laser-evoked firing rate response for all optotagged cells. Firing rates in **F** and **G** are normalized per neuron to peak = 1.

Significance was tested by comparing each neuron’s mean firing rate during the laser pulse to a null distribution generated by randomly shifting all laser onsets relative to the spike train 2,000 times. Each of the optotagged NPH neurons showed apparent AHV tuning and robust saccade-related activity, similar to the broader recorded population (Fig. 6E–F; Fig. S8). In particular, 4/5 optotagged neurons showed significant slow-phase modulation, and 5/5 showed significant quick-phase modulation. All five neurons also generated robust bursts of action potentials in response to the laser pulses (Fig. 6G). Per-neuron median latencies were within 10 ms of laser onset, averaging 4.40 ± 1.84 ms (mean ± SD of medians across *n* = 5 cells; range, 2.39–7.01 ms; corrected for TTL-to-shutter delay; see Methods). Thus, even anatomically defined NPH neurons projecting to DTN carried robust slow-and quick-phase eye-movement signals, arguing against a specialized projection-defined subpopulation that selectively relays AHV to the HD circuit.

### Modeling: NPH population activity supports flexible decoding of multiple orienting variables

Given that NPH neurons encode eye-movement signals that generalize across behavioral contexts rather than a dedicated angular head velocity (AHV) signal, we next asked whether accurate head-direction signals could nevertheless emerge from the structure of the population response. We therefore tested whether distinct downstream weightings of the experimentally observed NPH activity could reconstruct head direction despite its predominantly oculomotor structure. To address this, we built population models using the empirically measured relationships between slow-and quick-phase tuning. NPH neurons were classified into two dominant types: “same” neurons, in which slow-and quick-phase responses share directional preference, and “opposite” neurons, in which these components have opposing preferences (Fig. 7A). Using these experimentally constrained tuning properties, we simulated population activity during VOR-like behavior and examined how alternative downstream weighting schemes transformed the same activity into estimates of different orienting variables.

**Figure 7.**
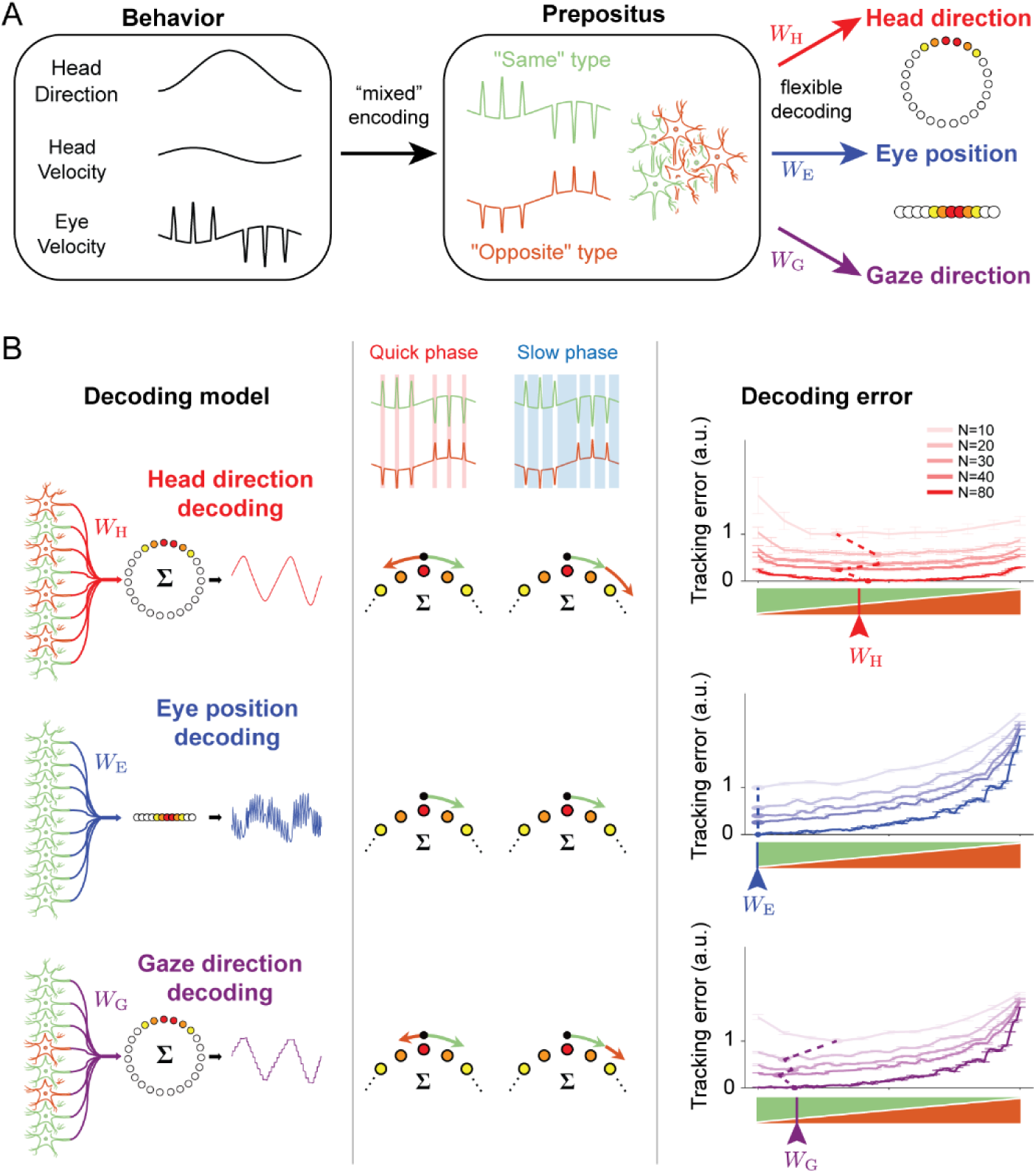
NPH population activity supports flexible decoding of head, eye, and gaze signals through structured population readout. **A**) Schematic of heterogeneous NPH population coding. Neurons are grouped into two experimentally defined classes: “same” neurons (green), in which slow-and quick-phase responses share directional preference, and “opposite” neurons (orange), in which these components have opposing preferences. This organization creates a mixed population code in which saccade-related (quick-phase) activity is distributed with opposing signs across the population. **B**) Decoding multiple orienting variables from the same NPH population depends on downstream readout structure. **Top**: Accurate head direction signals emerge only when inputs from “same” and “opposite” neurons are combined with weights that selectively cancel quick-phase activity while preserving slow-phase components. **Middle**: Readouts biased toward “same” neurons yield signals tracking eye position. **Bottom**: Unconstrained combinations of neuron types produce gaze-related signals that reflect both head-and eye-centered components. **Right**: Decoding error as a function of cell type ratio and population size. The x-axis represents the relative proportion of “same” and “opposite” type neurons; different colored lines correspond to different population sizes. Accurate head direction estimation requires a heterogeneous population and improves with increasing neuron number, reflecting averaging across heterogeneous responses. W_H_ = head direction decoding model weights. W_E_: eye position decoding model weights. W_G_: gaze direction decoding model weights.

**Figure 8.**
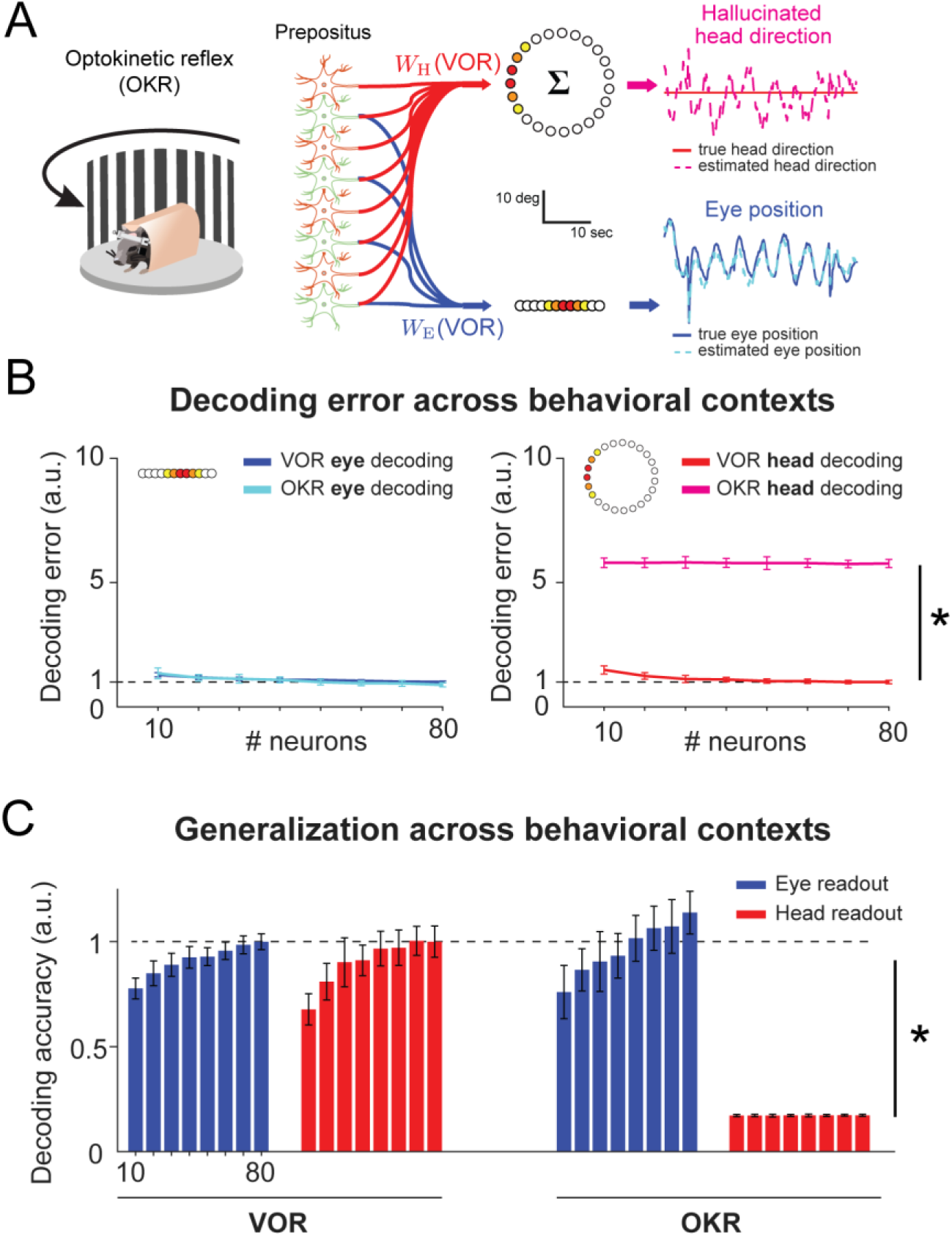
Fixed decoding weights reveal context-dependent failure of head direction estimation. **A**) Generalization test across behavioral contexts. Decoding weights for head direction and eye position were derived under vestibulo-ocular reflex (VOR) conditions (as in Fig. 7) and applied unchanged to population activity during optokinetic reflex (OKR), in which eye movements occur without head motion. Under VOR conditions, both head direction (red) and eye position (blue) are accurately reconstructed. Under OKR conditions, eye position decoding remains accurate (blue), whereas head direction decoding produces a spurious estimate of head motion (magenta), reflecting the misinterpretation of eye-movement signals as head velocity. **B**) Population-level decoding error (log scale) across behavioral contexts, as a function of population size. **Left**: Eye position estimation error (blue lines) is low under both VOR and OKR conditions. **Right**: Head direction estimation error is low under VOR conditions (red line) but markedly elevated under OKR conditions (magenta line). Increasing population size reduces variability but does not eliminate the systematic bias in head direction estimates under OKR. Dashed line indicates the normalized reference value, decoding error at N=80 neurons. **C**) Same data as in B), but here showing decoding accuracy (linear scale). Dashed line indicates the normalized reference value (1.0). The decoding model improves in accuracy for increasing population size. Eye position decoding accuracy is maintained across contexts, whereas head direction decoding fails to generalize from VOR to OKR.

Within the model, accurate head-direction estimates emerged only when “same” and “opposite” neurons were combined with structured weights that canceled saccade-related components while preserving the slow-phase drive associated with head rotation (Fig. 7B, top). In contrast, populations dominated by a single neuron type failed to produce accurate estimates: “same”-dominated populations generated large transient deviations driven by saccade-related activity, whereas “opposite”-dominated populations showed reduced but persistent deviations due to incomplete cancellation. Thus, stable head-direction estimates did not arise directly from the tuning of individual NPH neurons, but required a population-level readout that suppressed their dominant saccade-related signals.

Importantly, different weightings of the same NPH population activity reconstructed distinct orienting variables. Weighting schemes that emphasized “same” neurons yielded eye-position estimates, whereas alternative combinations of the two response classes reconstructed gaze direction (Fig. 7B, middle and bottom), where gaze is eye position plus head direction. The relative contribution of “same” and “opposite” neurons to each readout is shown in Fig. S10. Increasing population size further reduced decoding error, consistent with averaging across heterogeneous responses (Fig. 7B, right, shading, light to dark). Together, these results show that robust estimates of eye position, gaze direction, and head direction can all be reconstructed from the same NPH population activity, with the decoded variable determined by downstream weighting rather than by a dedicated AHV channel.

### Modeling: VOR-derived head-direction readouts fail when eye and head movements dissociate

Having established that accurate head-direction estimates can be reconstructed from NPH population activity through appropriate downstream weighting (Fig. 7), we next asked whether these readouts generalize across behavioral contexts in which eye and head movements become dissociated. Because NPH activity is dominated by eye-movement signals, a decoder trained during the vestibulo-ocular reflex (VOR), when eye and head movements are tightly coupled, could appear to recover head direction while actually reading out the correlated eye-movement component.

To test this, we derived decoding weights for head direction and eye position exclusively during simulated VOR and then held these weights fixed while applying them without refitting to simulated optokinetic reflex (OKR) activity, in which the eyes moved while the head remained stationary. Thus, any failure to generalize reflected a mismatch between the fixed readout and the decoded behavioral variable, rather than a change in the decoder itself. Under VOR conditions, both head direction and eye position were accurately reconstructed from the same population activity, consistent with the strong coupling between these variables (Fig. 8A,B). However, when the same fixed weights were applied to OKR data, a clear dissociation emerged. Eye position decoding generalized across contexts, continuing to accurately track eye movements (Fig. 8A, blue). In contrast, the head-direction readout failed systematically, generating large spurious (“hallucinated”) changes in estimated heading despite the complete absence of head movement (Fig. 8A, magenta).

At the population level, this dissociation was robust. Head-direction estimation error remained low under VOR conditions but increased dramatically under OKR (2-way ANOVA, context factor: F(1,784) = 1.72 × 10^5^, p <.001, η_p_² = 0.996), and this error persisted even as population size increased (population size factor: F(7,784) = 31.28, p <.001, η_p_² = 0.218, Fig. 8B). Increasing population size reduced variability but did not eliminate the systematic bias, indicating that the decoding failure reflected an incorrect mapping between neural activity and the inferred variable rather than noise in the representation. In contrast, eye-position decoding generalized robustly across contexts and improved with increasing population size (context factor: F(1,784) = 27.54, p <.001, η_p_² = 0.034; population size factor: F(7,784) = 132.67, p <.001, η_p_² = 0.542). Similar results were observed in the decoding accuracy (Fig. 8C).

Together, these results show that a head-direction readout learned when eye and head movements are coupled does not generalize when those variables dissociate. Fixed VOR-derived weights interpreted OKR-related eye-movement activity as head motion, producing erroneous updates of estimated heading despite a stationary head. By contrast, eye-position readouts remained accurate across contexts. Thus, the apparent success of decoding head direction from NPH activity during VOR does not establish that NPH carries a dedicated AHV signal; it can instead arise from the unmeasured coupling between eye and head movements.

## Discussion

### NPH carries predominantly oculomotor rather than dedicated AHV signals

Our findings overturn the prevailing model in which the nucleus prepositus hypoglossi (NPH) supplies a dedicated angular head velocity (AHV) signal to the mammalian head-direction system. Canonical attractor models posit that AHV-related inputs rotate the activity packet as the animal turns (Skaggs et al., 1995; Redish et al., 1996; Zhang, 1996; Song and Wang, 2005; Taube, 2007; Laurens and Angelaki, 2018). Yet simultaneous measurement and experimental dissociation of eye and head movements revealed that neurons meeting conventional criteria for AHV tuning predominantly carried oculomotor signals. Most showed robust modulation during both slow and quick phases of eye movement, and eye-movement sensitivities estimated in the absence of head motion predicted NPH activity during head rotation with little additional explanatory power from head velocity. Robust eye-movement signals were even present among anatomically identified NPH neurons projecting to DTN, extending the oculomotor signal into the ascending pathway itself. Finally, empirically constrained population models showed that different weightings of the same heterogeneous activity could reconstruct eye position, gaze direction, or head direction. Thus, a canonical input to the HD pathway is not a dedicated AHV signal, but a heterogeneous, predominantly oculomotor population code containing sufficient information for multiple possible orienting readouts.

The identification of NPH as an AHV relay rested not only on its anatomical inputs from the medial vestibular nucleus (MVN; McCrea, 1988) but also on two further observations that appeared mutually reinforcing. When NPH activity is analyzed solely against head kinematics, many neurons satisfy established criteria for AHV tuning, including the asymmetric and rectified response profiles predicted to update a ring attractor (McCrea et al., 1979; Graham et al., 2023). Inactivation of NPH also destabilizes HD tuning, demonstrating that this pathway is required for maintaining a stable directional representation (Butler and Taube, 2015; Butler et al., 2017). Together, these findings seemed to identify both the transmitted variable and its computational function. However, tuning and causal necessity do not establish signal identity when candidate behavioral variables covary. During the vestibulo-ocular reflex (VOR), compensatory slow-phase eye velocity is approximately equal and opposite to head velocity, including in mice (van Alphen et al., 2001; Stahl, 2004; Cullen and Taube, 2017). Eye-velocity coding can therefore produce compelling AHV tuning when eye movements are not measured. Earlier studies established that NPH activity covaried with head velocity and that NPH was necessary for HD stability, but they could not determine which component of coordinated eye–head behavior its neurons encoded.

Experimentally dissociating eye and head motion resolved this ambiguity at the level of NPH. Sensitivities estimated from eye movements produced without head motion predicted neural responses during VOR without refitting, whereas head velocity explained little additional variance. Neurons classified as clockwise or counterclockwise AHV cells under conventional analyses were frequently better characterized by oppositely directed slow-phase eye movements. Most also carried prominent quick-phase signals, ruling out the possibility that NPH output is simply an inverted replica of head velocity.

These results reconcile two literatures that had assigned incompatible computational roles to the same nucleus. Decades of anatomical, physiological, and lesion studies identify NPH as a core component of the premotor circuitry for horizontal eye movements and gaze stabilization (McCrea and Baker, 1985; Langer et al., 1986; Belknap and McCrea, 1988; Robinson et al., 1994; Iwasaki et al., 1999). NPH neurons encode eye position, eye velocity, and saccade-related drive during fixation, pursuit, and vestibular stimulation, while focal lesions impair gaze holding and both slow and quick phases of vestibular and optokinetic responses (Lopez-Barneo et al., 1982; Cheron et al., 1986a, 1986b; Cannon and Robinson, 1987; McFarland and Fuchs, 1992; Cullen et al., 1993; Escudero et al., 1996; Kaneko, 1997, 1999; Sylvestre et al., 2003; Dale and Cullen, 2013, 2015). This organization is conserved across vertebrates: the homologous area I in fish functions as a gaze-holding integrator composed largely of neurons with eye-movement-related activity (Aksay et al., 2000, 2003; Debowy and Baker, 2011; Miri et al., 2022). The oculomotor and HD literatures therefore appear to have interpreted largely overlapping, predominantly eye-movement-related population activity through different behavioral measurements.

The projection-defined recordings rule out the most direct alternative explanation: that the broader NPH population is oculomotor whereas a specialized subset projecting to DTN selectively carries AHV. All five optotagged DTN-projecting neurons showed significant quick-phase modulation, and four also showed significant slow-phase modulation. Eye-movement signals therefore enter the ascending NPH– DTN pathway itself and are not confined to NPH neurons projecting to established oculomotor targets. Because the projection-defined sample was small, it cannot establish the prevalence or full distribution of signals across DTN-projecting NPH neurons. Nevertheless, robust slow-and quick-phase modulation in every sampled projection neuron argues against a labeled-line account in which the NPH–DTN projection selectively relays AHV. The quick-phase signals are particularly consequential because resetting eye movements are an intrinsic and recurring component of eye–head coordination in both primates and mice, not occasional perturbations superimposed on an otherwise smooth head-velocity signal (Schiller and Tehovnik, 2015; Payne and Raymond, 2017; Itokazu et al., 2018; Meyer et al., 2020; Michaiel et al., 2020).

### Population readout does not establish signal identity

Our population modeling reframes the computational role of this heterogeneous input. Different weightings of the same NPH activity reconstructed eye position, gaze direction, or head direction, demonstrating that the information available from a population need not correspond to a single variable explicitly encoded by its individual neurons. In the modeled head-direction readout, stable estimates required combining neurons with different slow-and quick-phase response patterns, thereby suppressing saccade-related transients while preserving slower components correlated with head rotation. The model is not intended to specify the biological readout in DTN or LMN; rather it establishes that the same predominantly oculomotor population activity can support several distinct orienting computations. This places the NPH pathway within a broader framework of mixed selectivity and population coding, in which the same shared activity can support different computations according to the connectivity and demands of the receiving circuit (Rigotti et al., 2013; Fusi et al., 2016; Saxena and Cunningham, 2019).

Cross-context generalization provided a more stringent test of signal identity than decoding accuracy alone. During VOR, eye and head movements are tightly coupled, allowing fixed weights to reconstruct either eye position or head direction from the same NPH activity. When those weights were applied without refitting during optokinetic stimulation, where the eyes moved while the head remained stationary, the eye-position readout remained accurate, whereas the head-direction readout generated large spurious changes in estimated heading. This failure persisted as population size increased, demonstrating a systematic mismatch between the inferred variable and the neural activity rather than insufficient sampling or population noise. Successful decoding within one behavioral context therefore does not establish what a neural population represents; a decoder can exploit correlations among behavioral variables and accurately reconstruct one variable while primarily reading out another. In the case of NPH, the HD readout failed the more stringent test of accurate decoding across behavioral settings designed to break the correlations among behavioral variables. Thus, our simulations show that the observed NPH tuning alone is insufficient to support accurate head-direction updating across behavioral contexts, implying that downstream circuits must incorporate additional information and/or suppress saccade-related components.

### Reassessing the ascending head-direction pathway

These findings also force a reassessment of what successive stages of the ascending pathway encode. DTN and LMN have generally been interpreted within the same head-centered framework used to classify NPH neurons, often without simultaneous measurement of eye movements. Their activity may represent head direction or head velocity, but it may instead contain gaze-related signals, mixtures of eye and head variables, or intermediate representations that transform among eye-, head-, and world-centered coordinates. Mixed gaze and head representations in hippocampal, entorhinal, and retrosplenial circuits make these alternatives biologically plausible (Rolls and O’Mara, 1995; Georges-François et al., 1999; Killian et al., 2012; Meister and Buffalo, 2016; Jacob et al., 2017; Corrigan et al., 2023). NPH input may also be combined with vestibular, proprioceptive, visual, or motor-related signals distributed across additional pathways (Blair et al., 1998; Bassett and Taube, 2001; Sharp et al., 2001b; Yoder and Taube, 2014). Simultaneous recordings across NPH, DTN, and LMN during behaviors that independently vary eye, head, and gaze movements will therefore be essential—not merely to determine how an assumed HD update is constructed, but first to establish what variables these successive populations actually encode. Natural orienting will be especially informative because mice flexibly alternate between compensatory eye movements and coordinated eye–head gaze shifts (Sakatani and Isa, 2007; Meyer et al., 2020; Michaiel et al., 2020; Verdone et al., 2026).

More broadly, our results expose a fundamental problem in assigning computational functions to neural circuits. When behavioral variables are strongly correlated, tuning, causal necessity, and successful within-context decoding can converge on the same incorrect assignment of signal identity. In NPH, these approaches supported the influential model of a dedicated AHV relay, yet direct measurement and experimental dissociation revealed a predominantly oculomotor signal. This revision changes the computational problem that the HD system must solve: NPH does not supply the clean, labeled AHV channel assumed by canonical models, but a heterogeneous signal capable of supporting multiple orienting representations. Determining how directional codes are constructed will therefore require measuring and independently manipulating the relevant dimensions of eye, head, gaze, and body motion. More generally, neural computations cannot be identified from correlations observed within a restricted behavioral regime; they must survive experiments designed to break those correlations.

## Acknowledgements

Thanks to Kyle Smith for the loan of the laser used in the optotagging experiments, to Andrew Alvarenga and Kye Cooley for instrumentation; and Jeffrey Taube for discussion. This work was funded by National Institutes of Health 1U01-NS111695 (MvdM., K.Z., K.E.C.), NIH R01DC002390 (K.E.C.), and NIH R01DC018061 (K.E.C.).

## Author contributions

J.J.S. and M.v.d.M. conceived and designed the experiments in Experiment 1. H.H.V.C. and K.E.C. conceived and designed the experiments in Experiment 2. C.B., K.Z., and K.E.C. conceived and designed the computational modeling in Experiment 3. J.J.S. performed all experiments and analyses for Experiment 1, including the optogenetic identification experiments. H.H.V.C. performed all experiments and analyses for Experiment 2. C.B. and K.Z. performed the computational modeling. C.B. contributed to the analyses of the combined dataset. J.J.S., K.E.C., and M.v.d.M. wrote the manuscript with input from all authors. K.E.C. and M.v.d.M. jointly supervised the project.

## Ethics declaration

All animal procedures in Experiment 1 were approved by the Dartmouth College Institutional Animal Care and Use Committee (protocol 00002235). All procedures in Experiment 2 were conducted under approved animal protocols at Johns Hopkins University (protocol MO23M106).

## Data Availability Statement

The datasets generated during this study are not yet deposited in a public repository. The processed data necessary to reproduce the figures and statistical analyses, including spike times, eye-and head-movement time series, and cell classifications will be deposited in a public repository before publication and the access link/DOI will be added to the final manuscript. Supporting data required for editorial assessment or peer review are available from the corresponding authors upon request.

## Code availability

Custom code used for preprocessing, statistical analyses, decoding analyses, and population modeling will be deposited in a public GitHub repository before publication and the access link added to the final manuscript. Code required for editorial assessment or peer review is available from the corresponding authors upon request.

---

**Figure S1.**
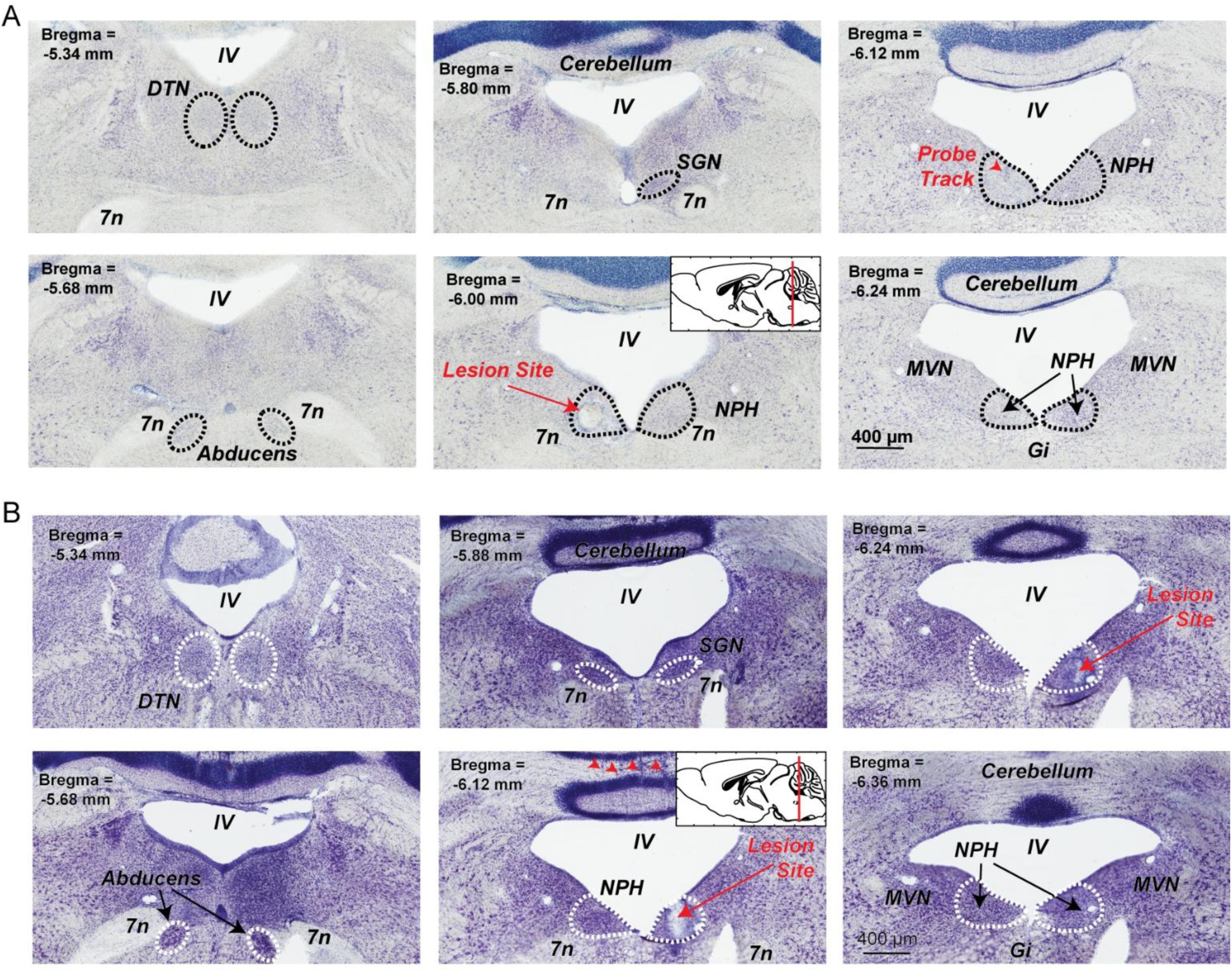
Example electrolytic lesions. Each panel shows a sequence of thionin-stained brain sections from a single mouse, running from anterior to posterior, column-wise. **A**) The electrolytic lesion site is indicated with a red arrow in the left NPH. **B**) Example histology and lesion site from a different mouse. Here, the electrolytic lesion site is in the right NPH. Isolated red arrowheads indicate silicon probe track marks. Abbreviations: DTN (dorsal tegmental nucleus), Gi (gigantocellular nucleus), IV (fourth ventricle), MVN (medial vestibular nucleus), NPH (nucleus prepositus), SGN (supragenual nucleus), 7n (facial nerve).

**Figure S2.**
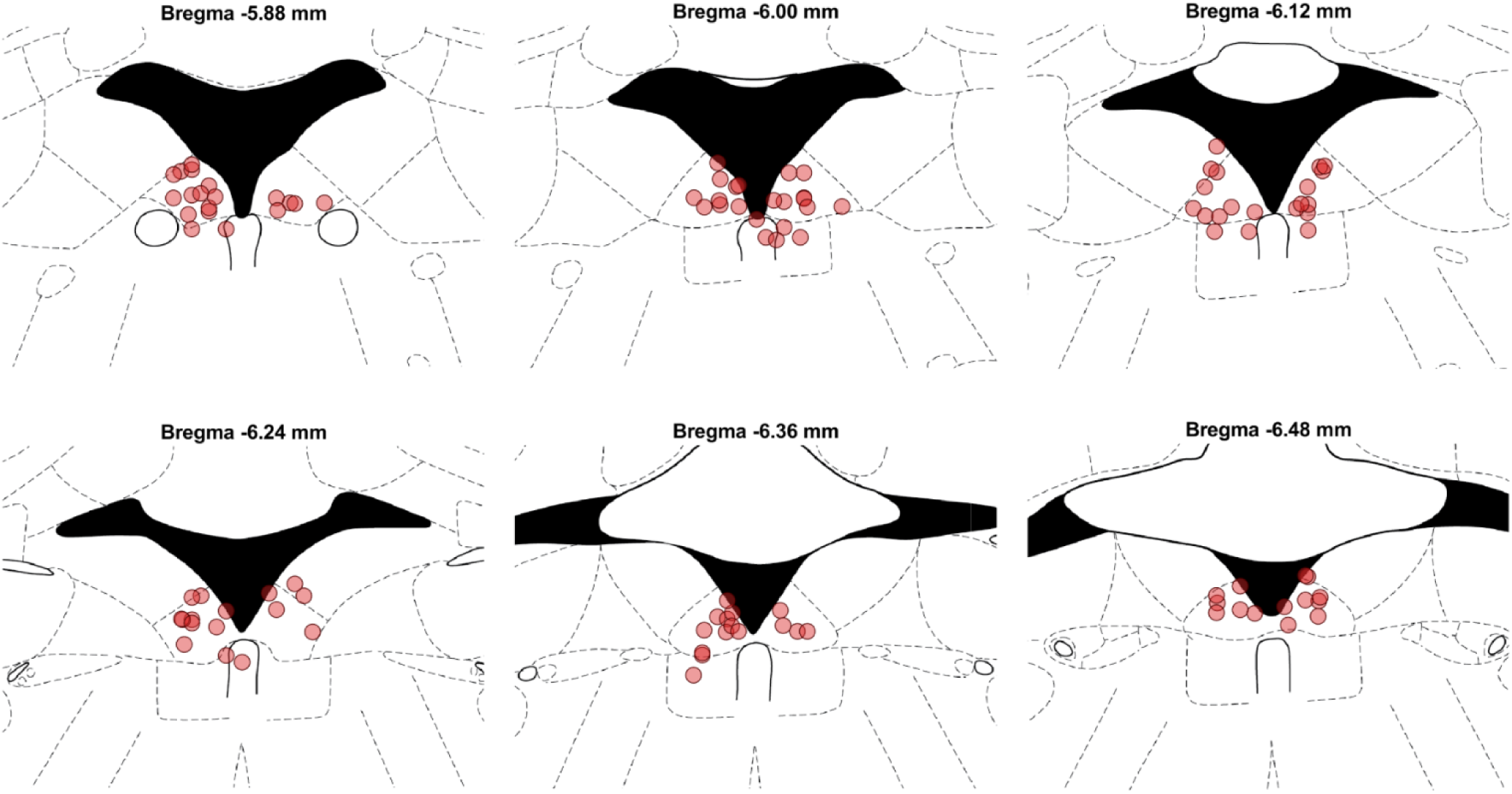
Depiction of recording locations. Each symbol indicates the location of a tetrode site in NPH. Recording site locations were determined by the presence of an electrolytic lesion mark or the terminus of a DiI probe tract in the *post mortem* tissue. For recordings with a lesion site or DiI track that had multiple good single units on separate tetrodes, those sites are included, based on the geometry of the probe.

**Figure S3.**
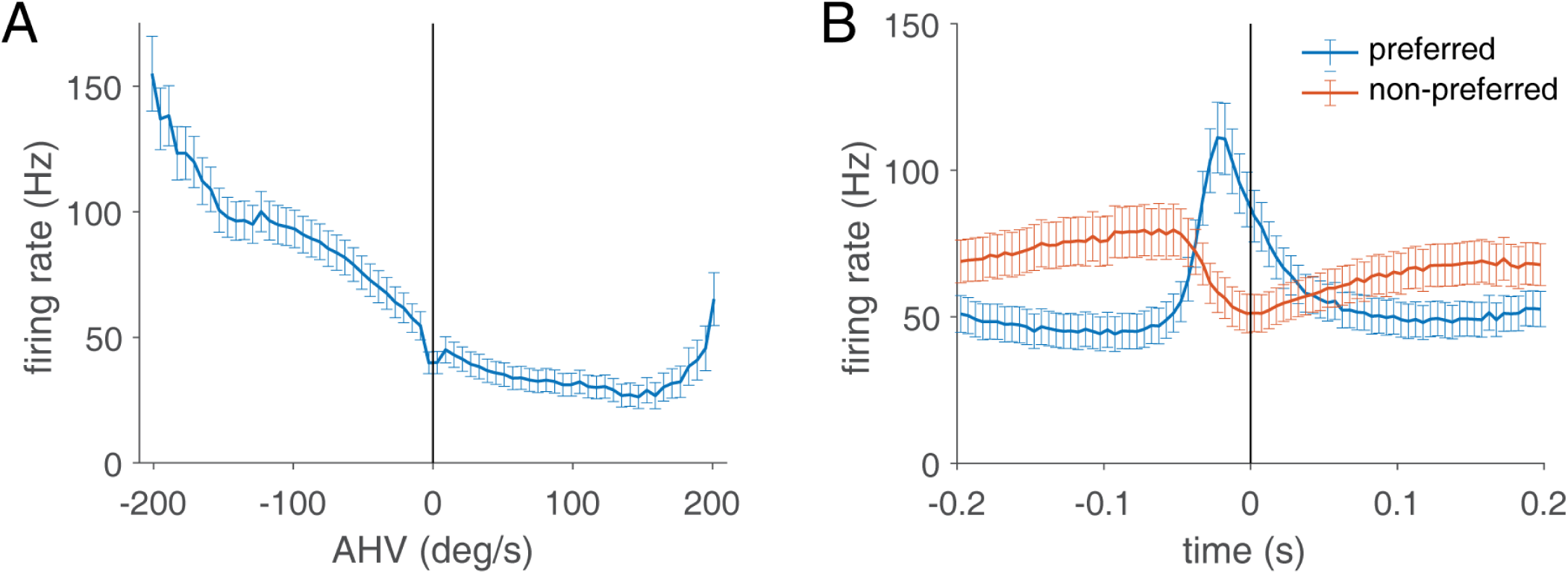
AHV tuning and saccade modulation are readily visible in the average firing rate response. **A**) The grand average tuning curve from all NPH neurons (*n* = 104) is shown, with each component tuning curve being oriented so that the preferred side has negative values. This is done for illustration purposes, to align the neurons. Error bars are S.E.M. The average of all AHV tuning curves is asymmetric, with a small dip at 0 deg, which corresponds to platform-stationary periods. **B)** Grand average PETH from all NPH neurons (*n* = 104), aligned to the time of saccade peak. Neurons are grouped by preferred and non-preferred saccade types. There is a clear burst of activity for preferred saccades and a significant dip in activity for non-preferred saccades.

**Figure S4.**
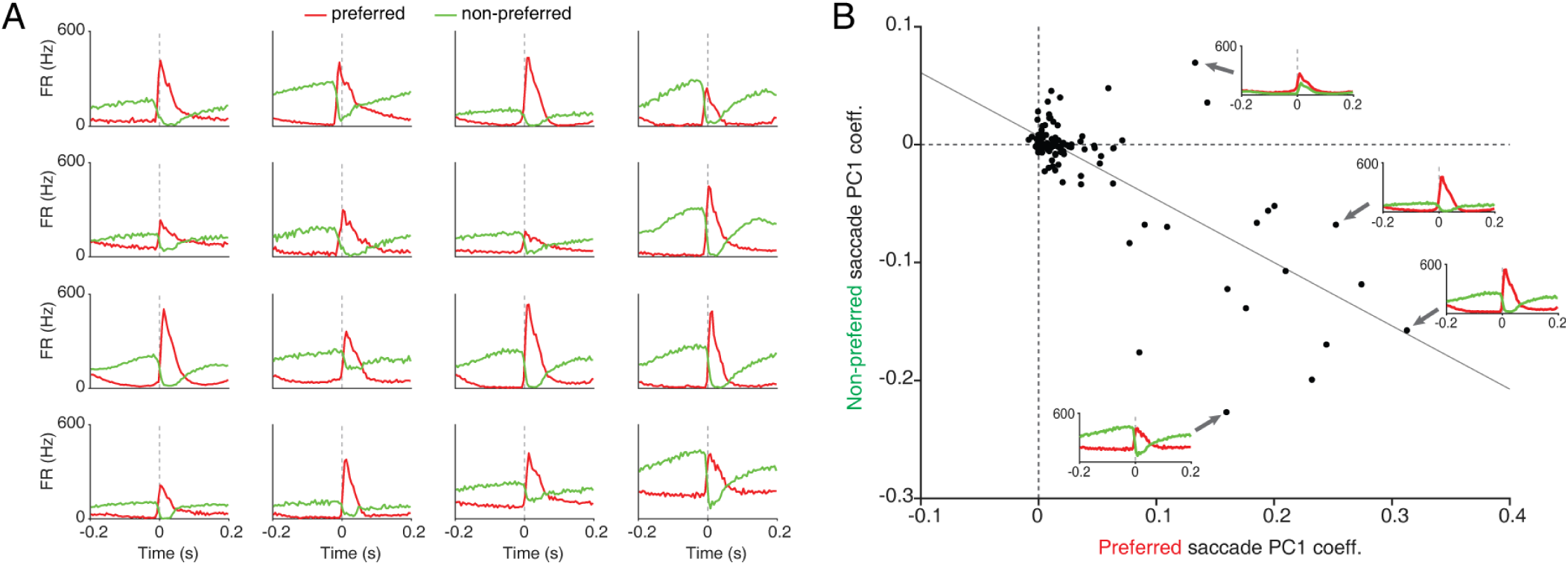
Quick-phase activity is bidirectional. **A**) Example peri-saccade firing rate histograms (PETHs) for 16 example NPH cells with strong quick-phase modulation. Red lines show firing rates for the preferred direction (i.e. nasal or temporal, depending on which direction is associated with a firing rate increase, see *Methods*) and non-preferred direction (green line); vertical line is the time of saccade onset. Note that peri-saccade firing rate changes tend to be opposite for the preferred and non-preferred directions. **B**) Scatterplot of principal component analysis (PCA) PC1 coefficients for the average preferred (red) and non-preferred (green) PETHs of each cell. Note that the line of best fit has negative slope, demonstrating opposite modulation for preferred and non-preferred saccades across the population of recorded neurons. Insets show the PETHs for individual examples (each data point in the scatterplot is one neuron).

**Figure S5.**
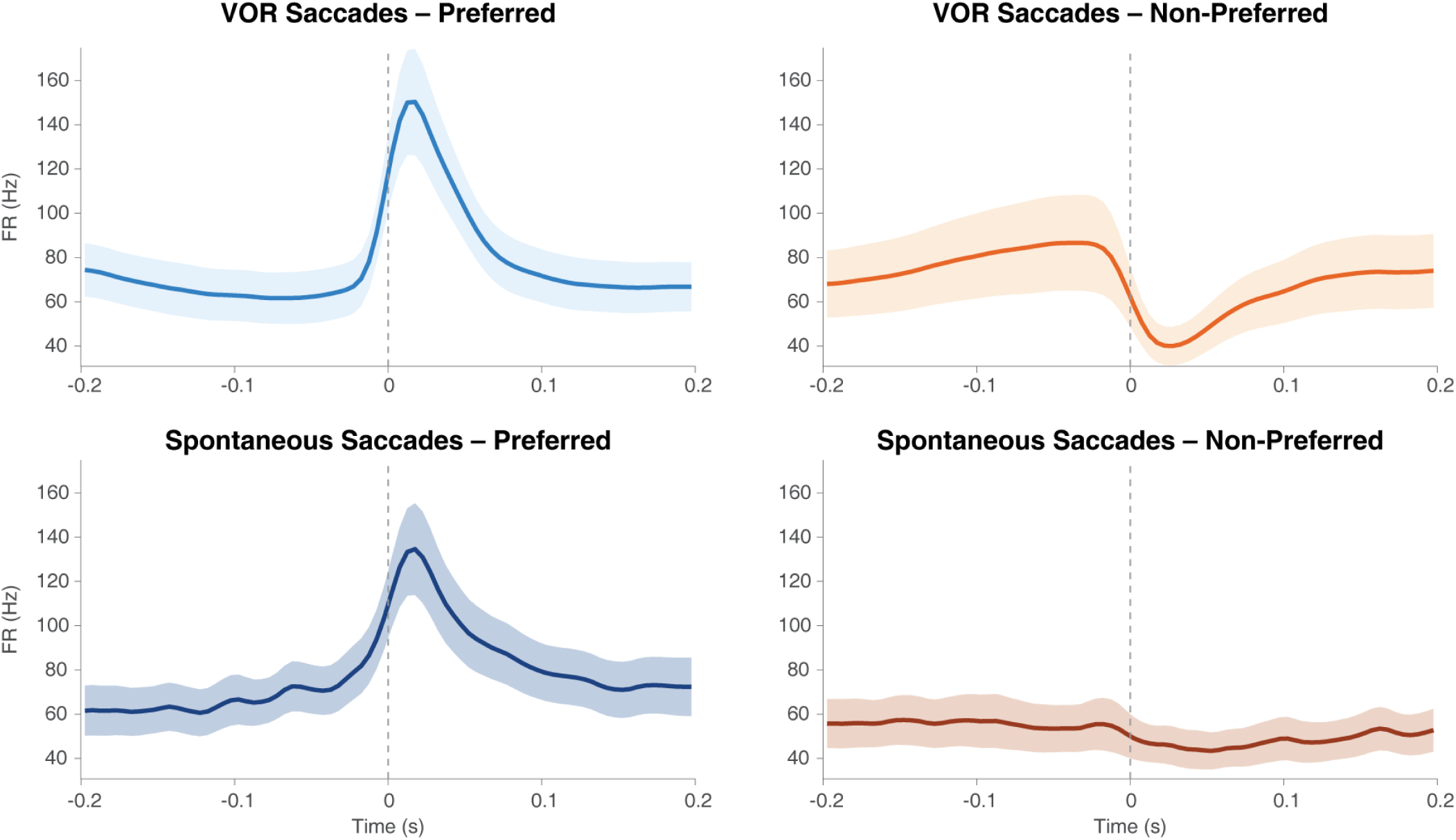
NPH saccade-evoked bursting persists during spontaneous saccades without head movement. Population peri-saccade time histograms (PETHs), aligned to saccade onset time, for 28 NPH neurons with ≥10 spontaneous saccades per direction, comparing VOR saccades (top row, platform moving) with spontaneous saccades (bottom row, platform stationary). Left column: preferred saccade direction; right column: non-preferred direction. Lines show population mean firing rate (Hz); shaded regions indicate ± SEM. Preferred direction was determined by the direction producing the larger burst during VOR saccades. The burst in the preferred direction is preserved during spontaneous saccades (Wilcoxon signed-rank test, VOR vs. spontaneous preferred burst: p = 0.96). These results demonstrate that NPH saccade-related bursting is a genuine oculomotor signal, not an artifact of vestibular input during the VOR.

**Figure S6.**
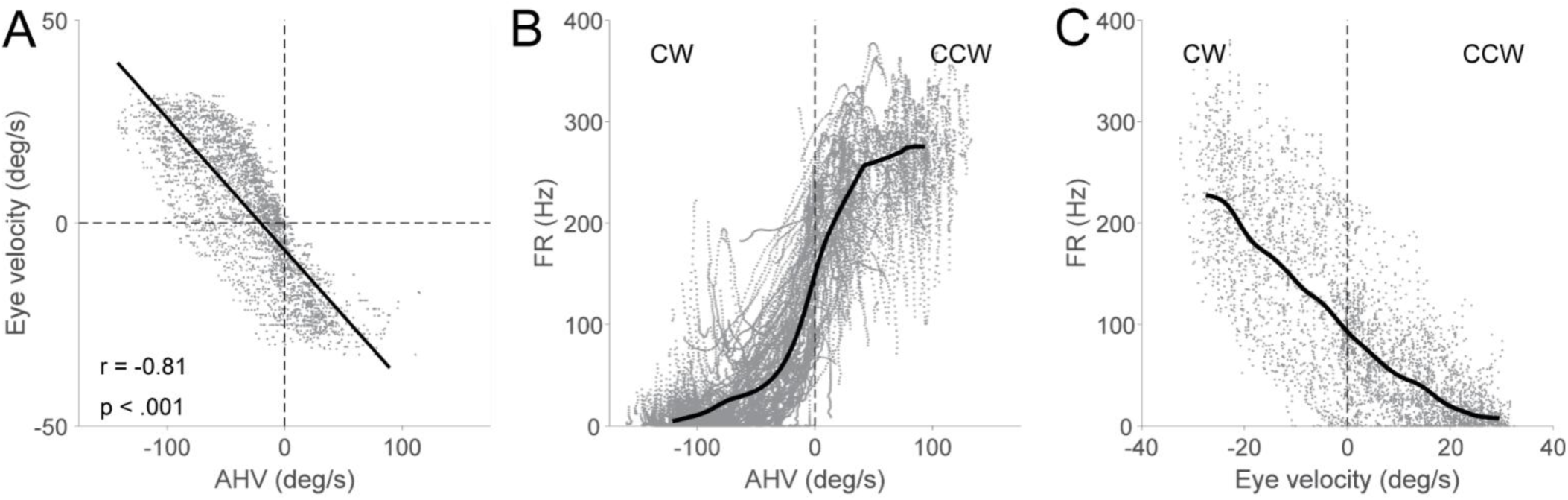
AHV and EV are inversely related, behaviorally and neurally. **A**) AHV and EV values are plotted for each time point for a single recording session (same session as in Fig. 3A). Note the inverse relationship and strong correlation, *r* =-0.81. **B**) AHV tuning curve for a single cell. This neuron “prefers” CCW rotation. **C**) EV tuning curve for the same neuron. In terms of EV tuning, this neuron prefers eye movements to the opposite direction (CW).

**Figure S7.**
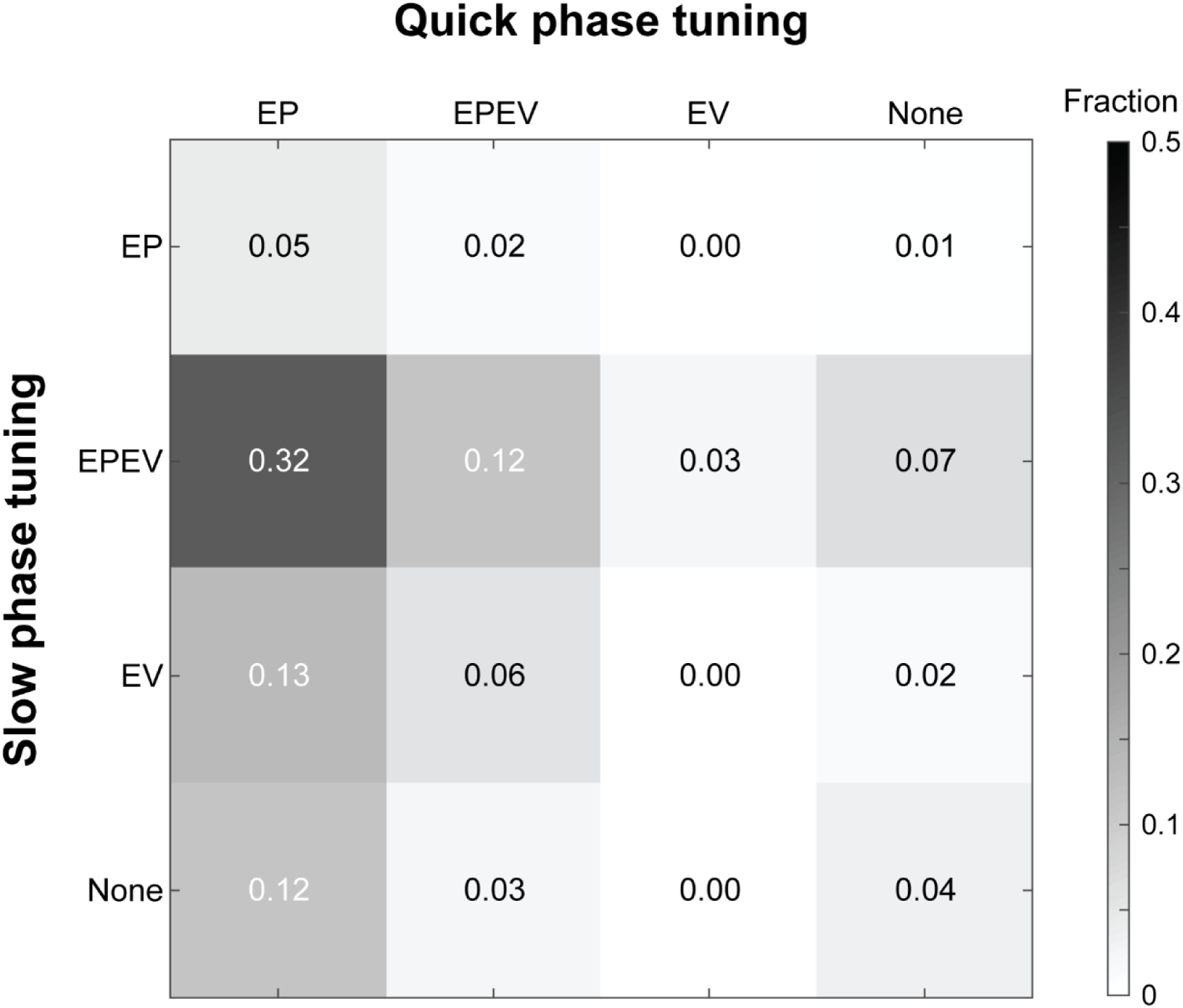
Generalized linear model analysis reveals subcategories of NPH neuron eye sensitivity. NPH neuron response to slow phase and quick phase can each be identified as both EP and EV sensitive, EP sensitive, EV sensitive, and none sensitive, based on GLM EP and EV fitting coefficient significance, in total 4 × 4 = 16 subcategories. The matrix shows the proportion in each subcategory. EP = eye position. EV = eye velocity. EPEV = eye position and eye velocity.

**Figure S8.**
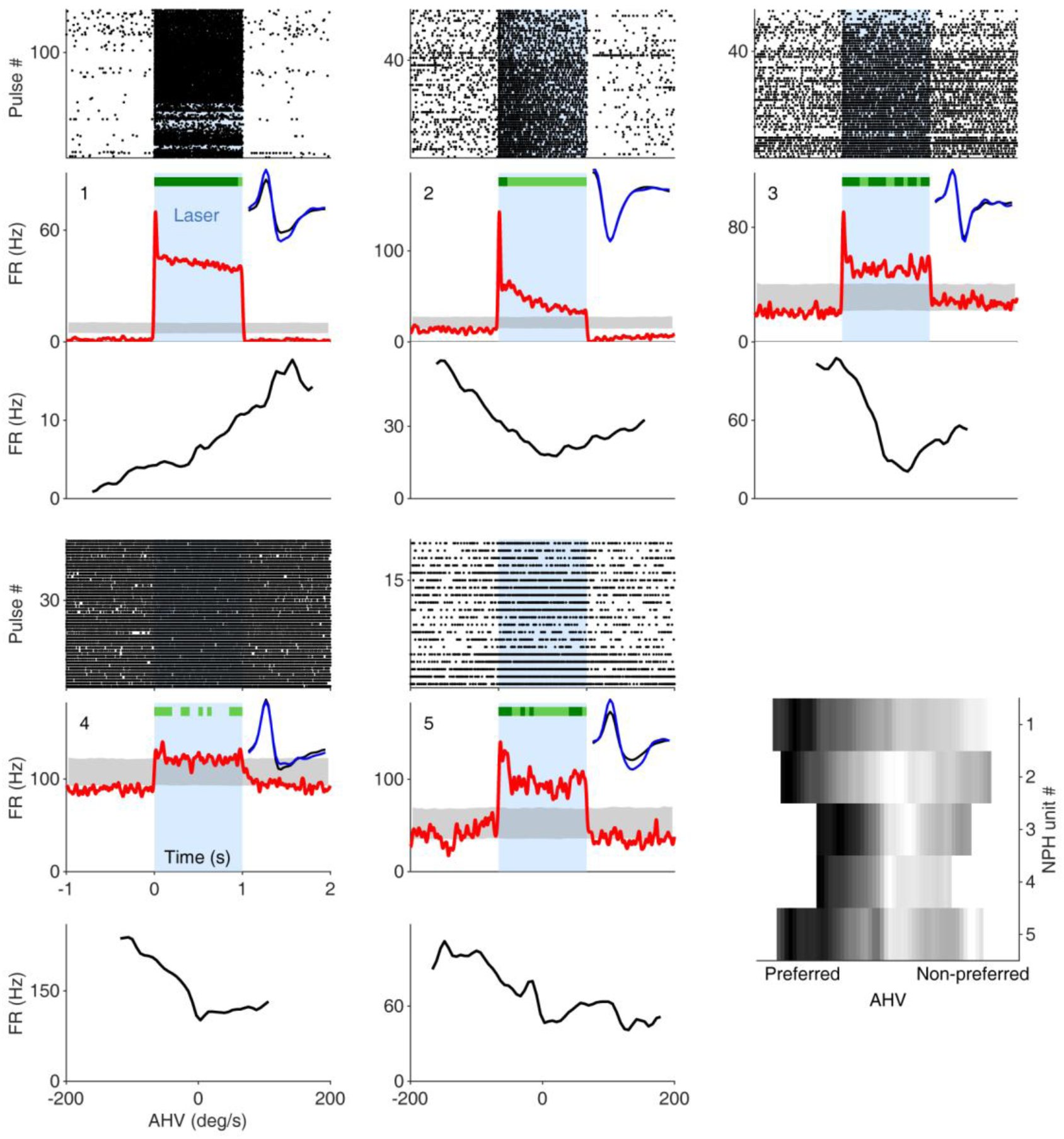
Optotagged cells. The first five panels each show data for one optotagged NPH→DTN neuron, organized as three vertically stacked sub-panels. **Top:** spike raster aligned to laser onset; black dots are individual spikes and blue shading marks the laser-on interval. **Middle:** peri-event time histogram (PETH) of firing rate aligned to laser onset. The red trace is the mean firing rate; the grey band shows the 5th–95th percentile envelope across 2,000 circular shuffles of the laser onset times. Dark green bars indicate per-bin significance at p <.001; light green bars indicate p <.05. **Inset (top right):** mean action-potential waveform on the peak channel for laser-evoked spikes (blue) and all other spikes (black). **Bottom:** angular head velocity (AHV) tuning curve for the same neuron. The sixth panel (bottom right) is a stacked heatmap of the AHV tuning curves for all five cells, aligned by preferred direction.

**Figure S9.**
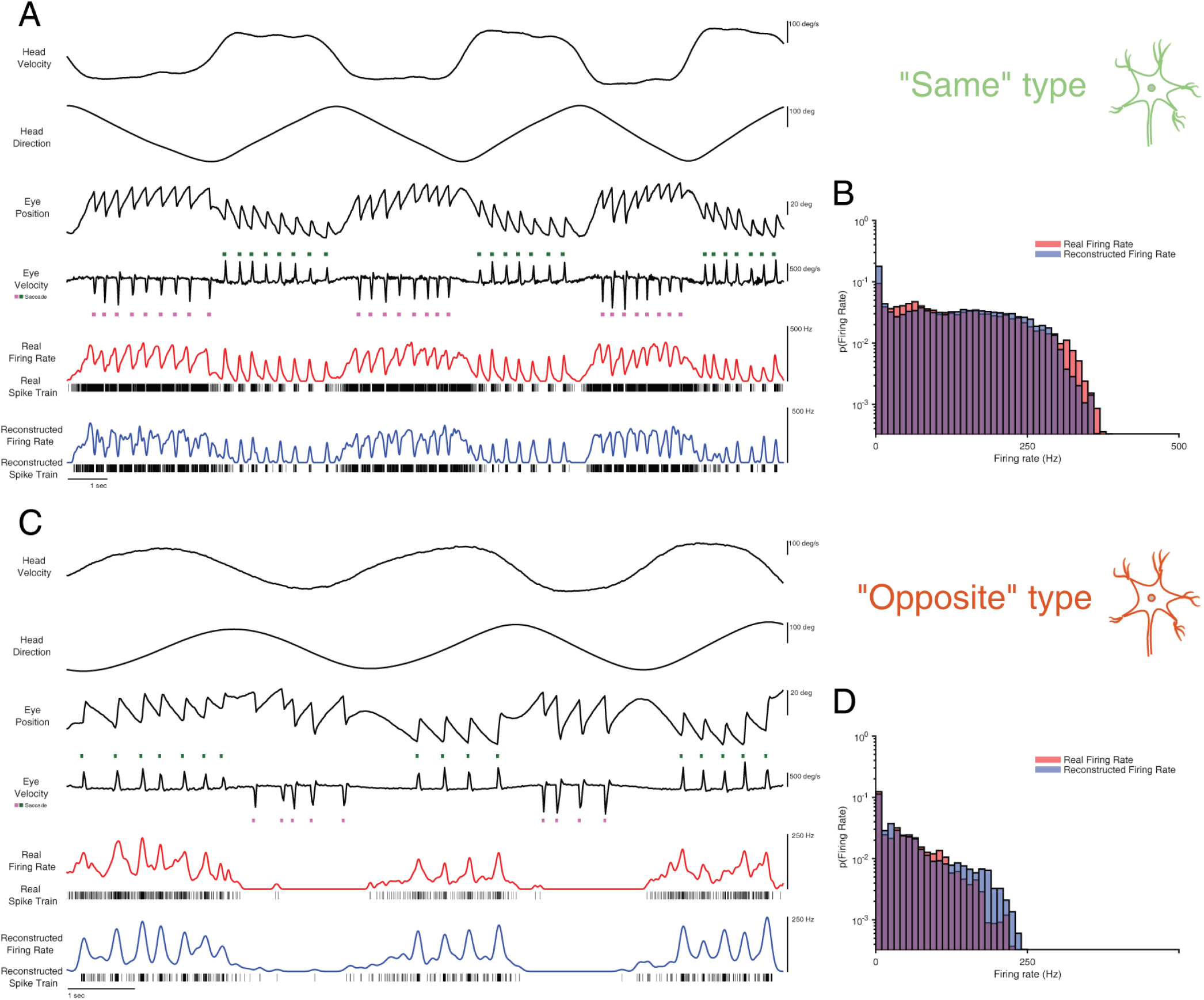
An eye-movement-based model reconstructs the firing rate of both “same” and “opposite” type NPH neurons during the VOR. **A)** Example “same” type neuron during sinusoidal passive head rotation. Traces from top: head velocity, head direction, eye position, and eye velocity. Saccade-onset ticks above and below the trace indicate saccade direction. The real firing rate (red) and spike train, and the reconstructed firing rate (blue) and spike train from the eye-based model (Methods). **B)** Probability distribution of the real (red) and reconstructed (blue) firing rates for the cell in (A); log y-axis. **C, D)** As in (A, B), for an example “opposite” type neuron. The distribution of reconstructed firing rates closely matches the real firing rate distribution for both cell types.

**Figure S10.**
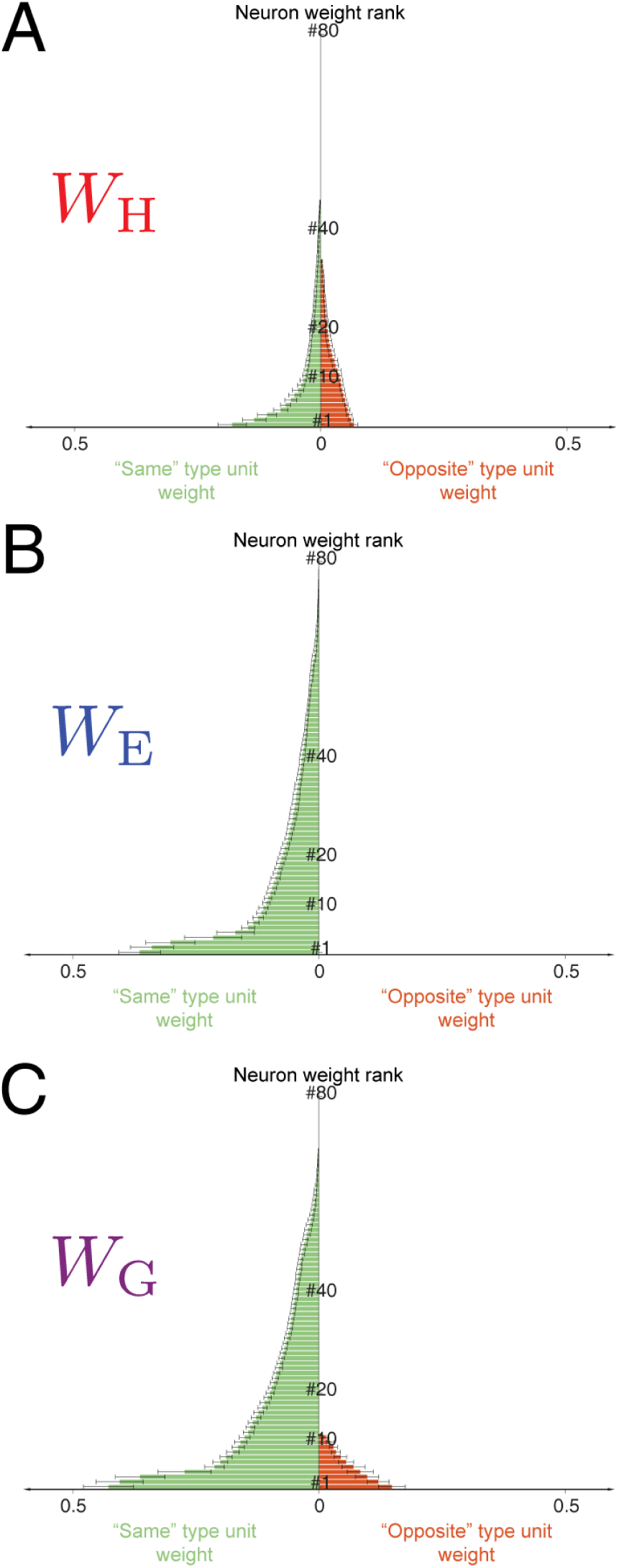
NPH unit weight distribution in the population decoding model. The “same” type (green) and “opposite” type (orange) NPH unit weight rank distribution in the optimal decoding model (Neuron number = 80) for **A**) head direction decoding (W_H_), **B**) eye position decoding (W_E)_, and **C**) gaze direction decoding (W_G_).

**Supplementary Table 1.** Breakdown of mouse sex, genotype, age, and number of sessions/neurons.

| Mouse | Sex | Strain | Age (mo) | Sessions | Neurons |
| --- | --- | --- | --- | --- | --- |
| M039 | M | C57BL/6J | 9 | 3 | 6 |
| M052 | M | C57BL/6J | 9 | 1 | 3 |
| M055 | M | C57BL/6J | 9 | 1 | 2 |
| M079 | M | C57BL/6J | 9 | 1 | 2 |
| M085 | M | C57BL/6J | 9 | 2 | 5 |
| M086 | M | C57BL/6J | 10 | 3 | 8 |
| M090 | M | C57BL/6J | 10 | 3 | 5 |
| M094 | M | C57BL/6J | 10 | 1 | 1 |
| M096 | M | C57BL/6J | 10 | 2 | 3 |
| M104 | M | C57BL/6J | 10 | 3 | 6 |
| M105 | M | C57BL/6J | 10 | 1 | 1 |
| M112 | M | C57BL/6J | 11 | 1 | 1 |
| M212 | M | C57BL/6J | 7 | 1 | 1 |
| M222 | F | CHETA | 13 | 2 | 3 |
| M247 | M | CHETA | 12 | 3 | 8 |
| M269 | M | Ai32 | 6 | 1 | 1 |
| M271 | M | Ai32 | 3 | 1 | 1 |
| M281 | M | Ai32 | 6 | 3 | 7 |
| M282 | M | Ai32 | 7 | 2 | 5 |
| M284 | M | Ai32 | 12 | 3 | 4 |
| M287 | F | Ai32 | 13 | 3 | 7 |
| M293 | M | Ai32 | 3 | 2 | 4 |
| M389 | F | Ai32 | 7 | 2 | 3 |
| M390 | M | Ai32 | 10 | 1 | 4 |
| M391 | M | Ai32 | 7 | 4 | 4 |
| M402 | M | Ai-27 | 5 | 3 | 7 |
| M444 | F | Ai32 | 5 | 1 | 2 |

